# Tear fluid as noninvasive liquid biopsy reveals proteins associated with malignant transformation of oral lesions

**DOI:** 10.1101/2025.11.06.686738

**Authors:** Erison S. Santos, Daniela C. Granato, Carolina M. Carnielli, Ariane F. B. Lopes, Daniella de Figueiredo, Luciana D. Trino, Fabio M. S Patroni, Bianca A. Pauletti, Romênia R. Domingues, Jamile Sa, Tatiane M. Rossi, Ana G.C. Normando, Narmeen Daher, Nataly Kravchenko-Balasha, Paul Debasis, Rosane Minghim, Luiz P. Kowalski, Alan R. Santos-Silva, Márcio A. Lopes, Thaís B. Brandão, Ana C. Prado-Ribeiro, Adriana F. Paes Leme

**Affiliations:** Laboratório Nacional de Biociências - LNBio, Centro Nacional de Pesquisa em Energia e Materiais - CNPEM, Campinas, SP 13083-100, Brazil; Departamento de Diagnóstico Oral, Faculdade de Odontologia de Piracicaba, Universidade Estadual de Campinas - UNICAMP, Piracicaba, SP 13414-903, Brazil; Institute of Dental Sciences, Faculty of Dental Medicine, The Hebrew University of Jerusalem, P.O.B. 12272, Ein Kerem, Jerusalem, ISRAEL 91120; School of Computer Science and Information Technology, University College Cork, Cork, Ireland; Head and Neck Surgery Department and LIM 28, University of São Paulo Medical School (FMUSP), São Paulo, Brazil; Department of Head and Neck Surgery and Otorhinolaryngology, A.C. Camargo Cancer Center, São Paulo, Brazil; Instituto do Câncer do Estado de São Paulo - ICESP, Faculdade de Medicina, Universidade de São Paulo - USP, São Paulo, SP 01246-000, Brazil

## Abstract

Oral leukoplakias (OLs) are premalignant lesions that can progress into oral squamous cell carcinoma (OSCC). This study hypothesized that tear fluid, as a noninvasive biofluid, reflects proteomic alterations associated with malignant transformation. The tear proteome of 44 individuals, including healthy controls, OL/PVL (proliferative verrucous leukoplakia), and OSCC patients, was deeply profiled, revealing 828 protein groups clustered according to histopathological alterations. N-glycoproteome analysis identified immune-related proteins, while public RNA-seq integration indicated immune imbalance marked by increased B-cell and decreased macrophage signatures during disease progression. Several immune-associated proteins and epithelial markers, including desmoplakin, KRT14, and DSC1, emerged as potential indicators of malignant transformation. These findings demonstrate that tear fluid reflects oral carcinogenic processes, thereby serving as a noninvasive liquid biopsy for early detection and clinical monitoring.

## INTRODUCTION

The poor prognosis of oral squamous cell carcinoma (OSCC) is associated with lymph node and distant metastases, which lead to recurrence, treatment resistance, and a low survival rate (1,2). This poor prognosis is partly due to late diagnosis, which often results in aggressive surgical treatment, causing facial deformities, reduced quality of life, and social isolation (1,2). Notably, 15.7% of patients experience suicidal ideation one year after diagnosis (3). OSCC can arise from oral leukoplakia (OL) and/or proliferative verrucous leukoplakia (PVL) (4). It is estimated that 9.5% of OL and 49.5% of PVL cases progress to OSCC (5). In addition, longitudinal follow-up studies indicate that the annual risk of malignant progression is 1.56% for OL and 9.3% for PVL (5). Comparatively, the annual progression rate of Barrett’s esophagus into adenocarcinoma is around 0.12% (6), highlighting the high malignant potential of OSCC. Currently, there is no therapeutic modality to interrupt this malignant transformation (7). Therefore, predicting the risk of OL/PVL progression to OSCC is crucial for early diagnosis and the identification of potential therapeutic targets.

In the clinical practice, the presence and grade of oral epithelial dysplasia are the main predictor of malignant transformation and histological analysis is used for patient risk stratification (8), which has many limitations. Firstly, there is considerable inter-pathologist variability in the assessment of the presence and grade of epithelial dysplasia (8). Secondly, even lesions without epithelial dysplasia can progress to OSCC (8). For example, the presence of this feature is not commonly observed in PVL, despite these lesions having a high risk of progression and recurrence (8-10). Because of this, patients undergo numerous invasive and uncomfortable biopsies over the years to evaluate changes in the tissue. However, lesions in the early stages promote minimal tissue alterations, and microscopical features may fail to predict malignant transformation. At these stages, imbalances in body fluid analytes may reflect the general pathophysiological status, favoring the discovery of molecular signatures. Therefore, noninvasive tools for identifying biomarkers of malignant progression would be valuable.

Advances in mass spectrometry-based proteomics have driving the discovery of biomarkers through noninvasive liquid biopsy (11). In the oral cavity, saliva is recognized as a promising type of liquid biopsy due to its proximity to oral lesions, which could provide advantages in biomarker discovery (12). However, translating saliva-derived biomarkers into clinical practice faces several challenges (13,14). For instance, elderly patients and those taking multiple medications have alterations in the amount and composition of saliva (15). In general, patients with OL, PVL, and OSCC are usually above 60 years old, and may be undergoing treatment for different diseases (7,16). Thus, identifying new liquid biopsy matrices beyond saliva is crucial to expand diagnostic possibilities.

In this context, noninvasive liquid biopsy of tear fluid represents a promising source of molecular signatures of oral diseases. It has been proposed as a source of biomarkers for ocular (dry eye, glaucoma, blepharitis) and systemic diseases (multiple sclerosis, diabetes), including cancers (breast, colon, ovary, lung) (17-20). Ocular and systemic diseases release circulating extracellular vesicles (EVs) carrying DNA, RNA, and proteins that modify the composition of tear fluid and indicate pathological processes occurring distant from the ocular surface (16-20). EV-derived proteins annotated from 37 different tissues, including tongue, were recently identified in this fluid (21). Despite its small volume, tear fluid contains low levels of albumin, does not require additional sample filtration or centrifugation as in plasma samples (22). Therefore, we hypothesized that discovery-based proteomics of tear fluid has the potential to reveal molecular signatures for oral malignant progression.

Hence, we employed a mass spectrometry-based proteomics approach to identify dysregulated proteins in different oral lesions that could predict malignant transformation. Taken together, our findings demonstrate that liquid biopsy reflects oral malignant progression and holds potential to identify biomarkers with clinical value.

## RESULTS

### Tear fluid Global proteome and *N*-glycoproteome are different across clinical groups

Unstimulated tear fluid samples from 44 individuals were collected from patients with oral leukoplakia (OL, N=10), with proliferative verrucous leukoplakia (PVL, N=10), with oral cancer (OSCC, N= 14) and from healthy controls (N=10) (table S1), and the global proteome and its respective non-enriched *N*-glycoproteome were obtained (Fig. 1A) (23). The statistically significant proteins and *N*-glycopeptides were associated with clinical and pathological data that may play a biological role in OL/PVL transformation to OSCC.

**Figure 1.**
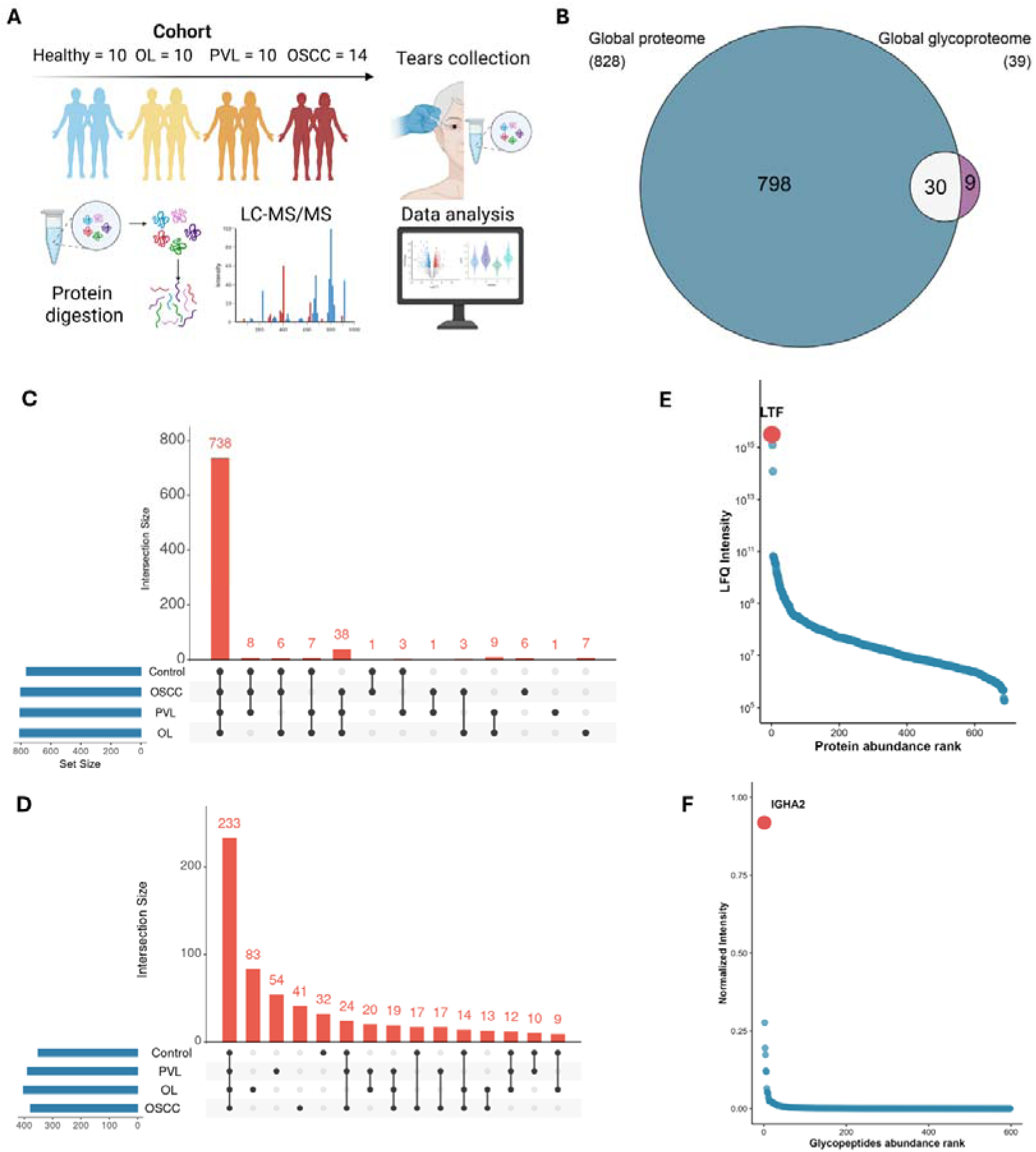
Characterization of global proteome and *N*-glycoproteome. **(A)** Experimental design to search for molecular signatures of malignant transformation in tear fluid. The cohort comprised patients with OL (n = 10), PVL (n = 10), OSCC (n = 14), and healthy controls (n = 10). Sixty microliters (µL) of saline solution were added to the ocular surface of each patient. Tear fluid was collected using a microcapillary glass tube and stored in Eppendorf tubes. Samples were digested with trypsin and analyzed by liquid chromatography–tandem mass spectrometry (LC-MS/MS). MaxQuant and Perseus were used to identify and quantify proteins, while Byonic was used to identify *N*-glycopeptides. **(B)** Venn diagram of common and unique proteins and *N*-glycoproteins identified in the proteome and *N*-glycoproteome searches. **(C)** Shared and ‘exclusive’ identified proteins among the samples analyzed. **(D)** Shared and ‘exclusive’ identified N-glycopeptides among the analyzed samples. **(E)** Dynamic range showing the relative abundance of proteins. **(F)** Dynamic range showing the relative abundance of N-glycopeptides. In this analysis, unique glycopeptide abundances were normalized by dividing each glycopeptide’s intensity by the total intensity sum of its sample, correcting for inter-sample variation.

The quality control metrics indicated reproducibility of label-free protein quantification (Pearson correlation coefficients; Fig. S1), with similar intensities and retention times for the three selected iRT peptides—SSAAPPPPPR (iRT_Peptide_1), GISNEGQNASIK (iRT_Peptide_2), and HVLTSIGEK (iRT_Peptide_3), and the trypsin autolysis peptides at m/z 421.7584 (2+), 523.2855 (2+), and 1106.0557 (2+) across the 44 runs (24) (Fig. S2). In the global proteome, 854 protein groups (PGs) were identified, with 828 remaining after excluding “only by site” entries and reverse sequences and requiring at least one valid value in one group (Fig. 1B and table S2). Most proteins (737 PGs) were consistently shared among the four clinical groups (control, OL, PVL, OSCC) (Fig. 1C).

598 N-glycopeptides from 39 proteins were identified, 30 of which were also detected in the global proteome (Fig. 1B and table S3). Among these, 233 were shared across control, OL, PVL, and OSCC (Fig. 1D). Notably, most N-glycopeptides (90%) carried complex or hybrid N-glycans, corroborating previous reports (25) (fig. S3). The dynamic range of the proteome and N-glycoproteome spanned approximately five and four orders of magnitude under normalized intensity, respectively. Lactotransferrin (LTF) and immunoglobulin heavy constant alpha 2 (IGHA2) showed high abundance (26,27) (Fig. 1E-F), whereas, as expected, albumin exhibited low abundance.

We next investigated whether global proteome and *N*-glycoproteome data could distinguish patients according to clinical grouping and clinicopathological features. LFQ intensities (proteome) and normalized intensities (N-glycoproteome) were analyzed by unsupervised hierarchical clustering using Kendall distance with Ward linkage. Clustering did not perfectly segregate patients by clinical group (Fig. 2A–B), but two major clusters were observed. We next used the proteins grouped in each cluster to perform biological process enrichment analysis, which revealed high enrichment of fatty acid biosynthesis, inflammation, and immune response (Fig. 2C and table S4). Notably, the marked immune enrichment observed in both N-glycoproteome clusters underscores the role of protein glycosylation in antigen recognition, a process essential for ocular surface protection.

**Figure 2.**
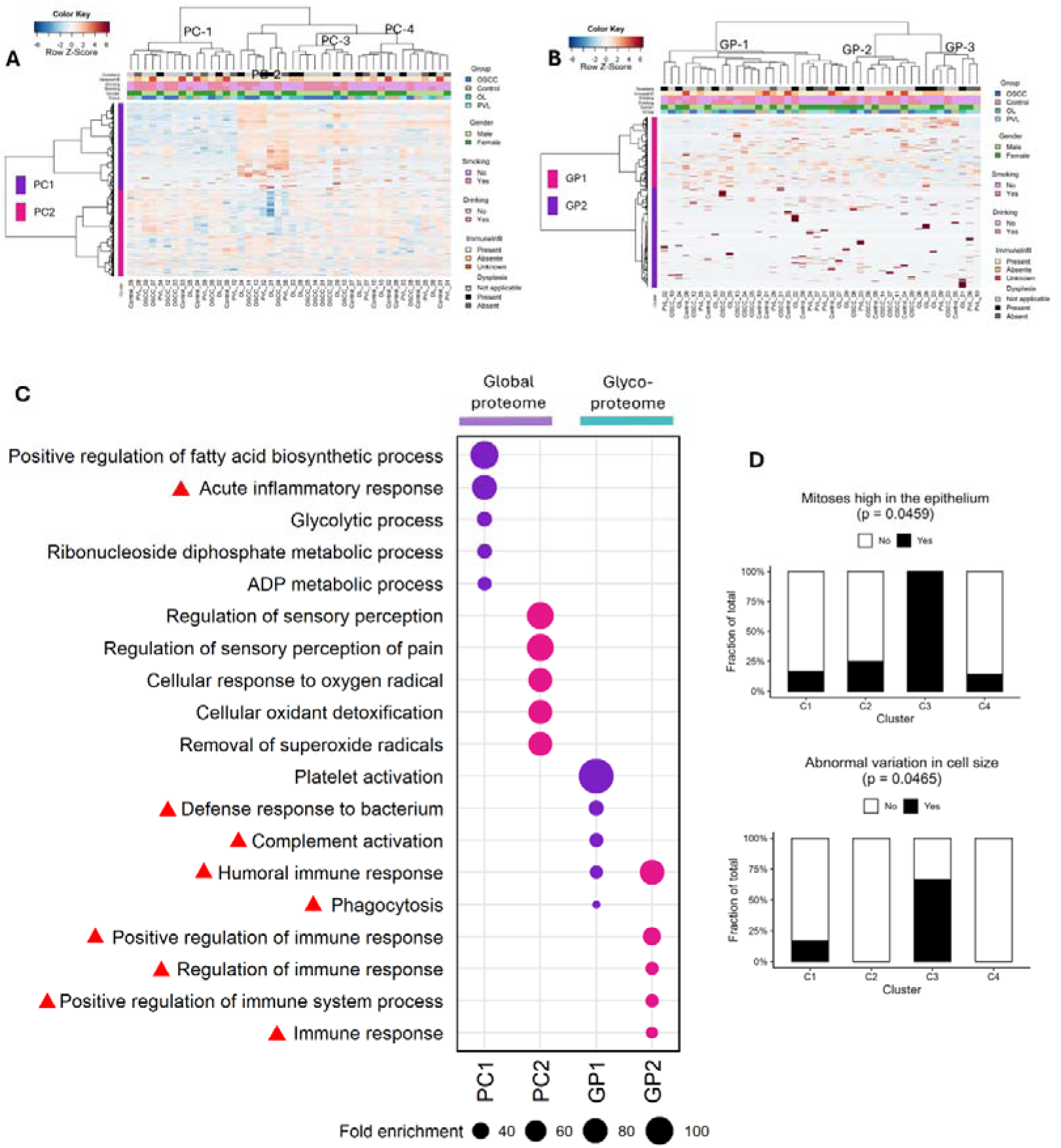
Global proteome and *N*-glycoproteome provides biological insights into oral diseases. **(A-B)** LFQ intensity values and normalized intensity values were used for unsupervised hierarchical clustering of the global proteome (n = 44) and N-glycoproteome (n = 44), respectively. Both analyses were performed using the *heatmap.3* function in the R environment with Kendall distance and Ward linkage. **(C)** Top five GO biological processes enriched in the protein clusters (PC1/PC2) and *N*-glycoprotein clusters (GP1/GP2) of the global proteome and *N*-glycoproteome, respectively (adjusted p ≤ 0.05; two-sided Fisher’s exact test followed by Benjamini–Hochberg correction). Immune-related biological processes are labeled with a red triangle. **(D)** Clinicopathological features that were significantly associated with patient clusters C1, C2, C3, and C4 in proteomic data (two-sided Fisher’s exact test; P≤ 0.05). These microscopic features are specific to patients with OL/PVL.

The global proteome and N-glycoproteome data separated the patients into four and three clusters, respectively. We next asked whether the clinical and pathological features of patients in these clusters differed significantly between clusters. This analysis demonstrated that two histological features assessed in patients with OL/PVL (mitosis in the epithelium and abnormal variation in cell size) differed significantly among patients according to proteomic data, but not in the N-glycoproteome data (Fig. 2D). These histological features were previously associated with malignant transformation (28,29). These results suggest that the global proteome of tear fluid can provide insight into malignant progression.

### Global proteome of tears indicates immune imbalance during malignant transformation

Considering that half of enriched biological processes on the global proteome were immune-related (Fig. 2C), we estimated the proportion of immune cells in our bulk proteome data by computational deconvolution. For this, we used CIBERSORTx (30) to infer the immune cell population present in the tear fluid based on our global proteomic data using a public dataset as reference. Previously published single-cell RNA sequencing (scRNAseq) data from normal tissue adjacent to OSCC tumor (N = 3), OL (N = 4), and OSCC (N = 10) were used as a reference for computational deconvolution (31). For this analysis, we included only OSCC HPV negative. Notably, tissue-derived data were used because there was no available public data of body fluids focusing on OL/PVL. In healthy samples, both the deconvoluted and the original tissue datasets showed a high fraction of fibroblasts. For OSCC samples, macrophages were highly abundant in the tissue data but showed low enrichment in the deconvoluted biofluid data (Fig. 3A–B). Overall, the deconvoluted tear fluid data displayed a high proportion of plasma B cells in OL, PVL, and OSCC (Fig. 3A). Interestingly, a high enrichment of B cells in lesions progressing toward malignancy has also been reported in laryngeal leukoplakia at the single-cell RNA level (32). These findings indicate that our deconvolution strategy is consistent with previous data and effectively captures the immune microenvironment of leukoplakias in the head and neck.

**Figure 3.**
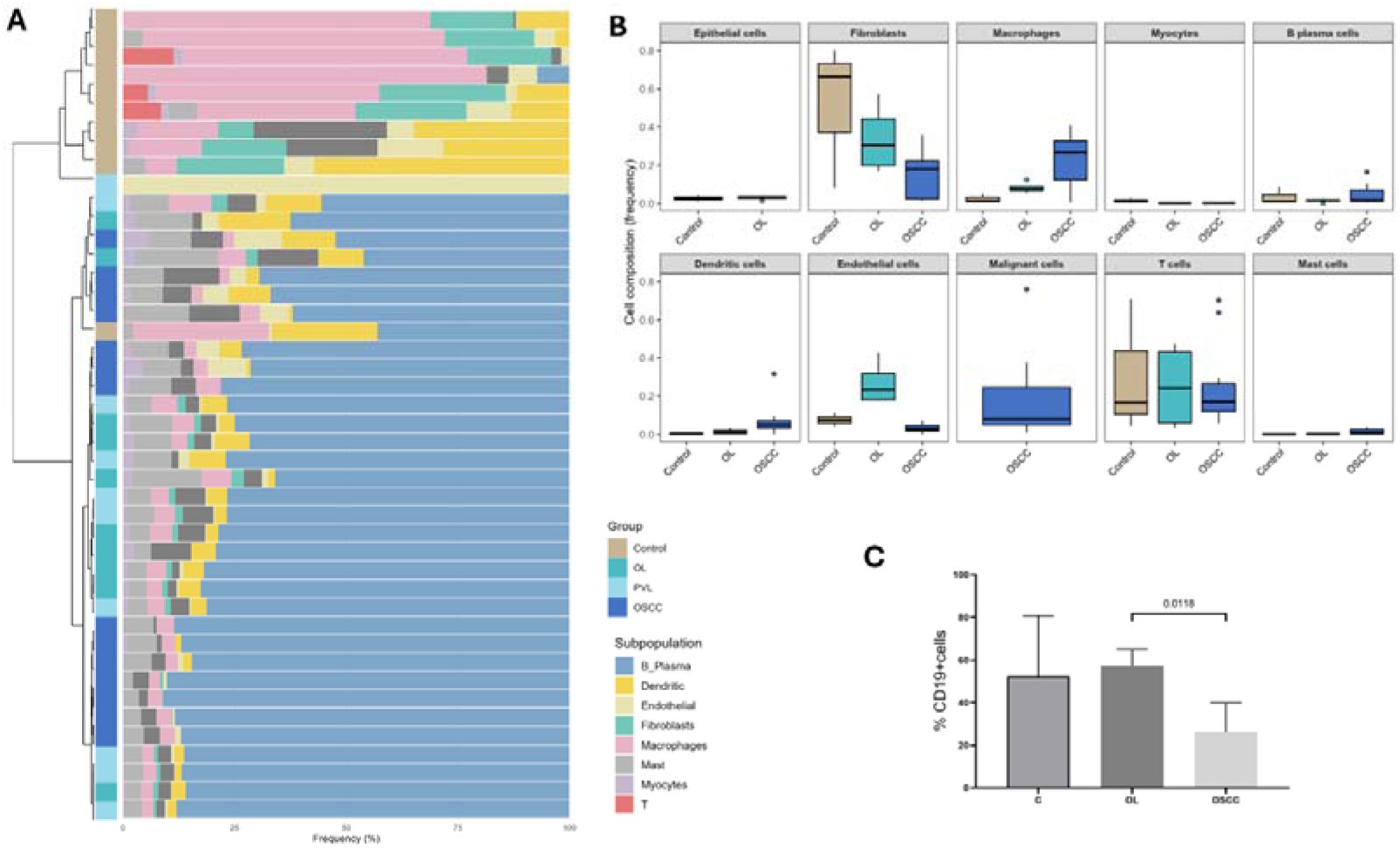
Immune cell composition of bulk tear fluid proteome according with computational deconvolution and flow cytometry analysis. **(A)** Relative frequency of cell composition from referenced scRNAseq from the tissue. Boxplots show the relative frequency of epithelial cells, fibroblasts, macrophages, myocytes, plasma B cells, dendritic cells, endothelial cells, malignant cells, T cells, and mast cells in healthy control, OL, and OSCC samples. Data points represent the average fraction of each cell type per sample, normalized to the total number of cells in that sample. Boxplots display the median (center line), interquartile range (box), and range (whiskers), with individual outliers shown as dots. **(B)** Prediction of cell populations based on the bulk proteome of tear fluid using CIBERSORTx version 1.0. The dendrogram was built using Euclidean distance and Ward linkage. **(C)** Percentage of total cells labeled as CD19 positive in controls (C), OL, and OSCC in flow cytometry analysis. It was performed one-way ANOVA followed by Tukey’s post hoc test; P ≤ 0.05 to compare the groups.

We further interrogated whether the proportion of deconvoluted immune cells could segregate patients according to clinical groups. Our hierarchical clustering analysis demonstrated that healthy control samples were completely segregated from OL/PVL and OSCC (Fig. 3A). Notably, the proportion of B cells in healthy control samples was lower compared to OL, PVL, and OSCC reinforcing that, as aforementioned, B plasma cells are abundant in these lesions. To obtain a deeper insight into B cells in tear fluid, we collected samples from an independent cohort of 32 patients and performed flow cytometry. This cohort comprised 7 controls, 9 OL, and 16 OSCC patients. These samples were pooled then analyzed. B cells decreased significantly in OSCC compared to OL but not compared to healthy controls (Fig. 3C). Lastly, we performed a Spearman correlation analysis to verify whether the proportion of deconvoluted immune cells were correlated with clinicopathological features. In OSCC samples, a positive correlation was observed between the presence of angiolymphatic invasion and the fractions of macrophages. In OL/PVL samples, a negative correlation was observed between the abundance of dendritic cells and high mitotic activity in the epithelium (table S5).

### Dysregulated proteins of tears reflect clinical and pathological features of oral malignant transformation

We demonstrated that the global proteome was able to provide insights into the malignant transformation process and distinguish patients according to their clinicopathological profile. We therefore hypothesized that tear fluid could present proteins differentially abundant across the clinical groups. To test it, we first performed pairwise two-sided comparisons of each clinical condition (unpaired Student’s t□tests or Welch’s t□tests followed by Benjamini–Hochberg correction; P□≤□0.05). Next, we carried out differential abundance analysis of all clinical groups (one□way ANOVA followed by Tukey’s post□hoc test; P□≤□0.05). We identified proteins that were up or downregulated in each pairwise comparison, except for PVL vs OSCC (Fig. 4A, and table S6). Also, we identified 40 differentially abundant proteins across all groups, five of which were also differentially abundant in N-glycoproteome (Fig. 4B, and table S7). Using previous public data employed in the computational deconvolution (31), we investigated which immune single-cells these differentially abundant proteins from the tear fluid were being expressed at the tissue level. Interestingly, some proteins were differentially expressed in immune cells at tissue level, particularly macrophages, T cells, and B cells (Fig. 4C).

**Figure 4.**
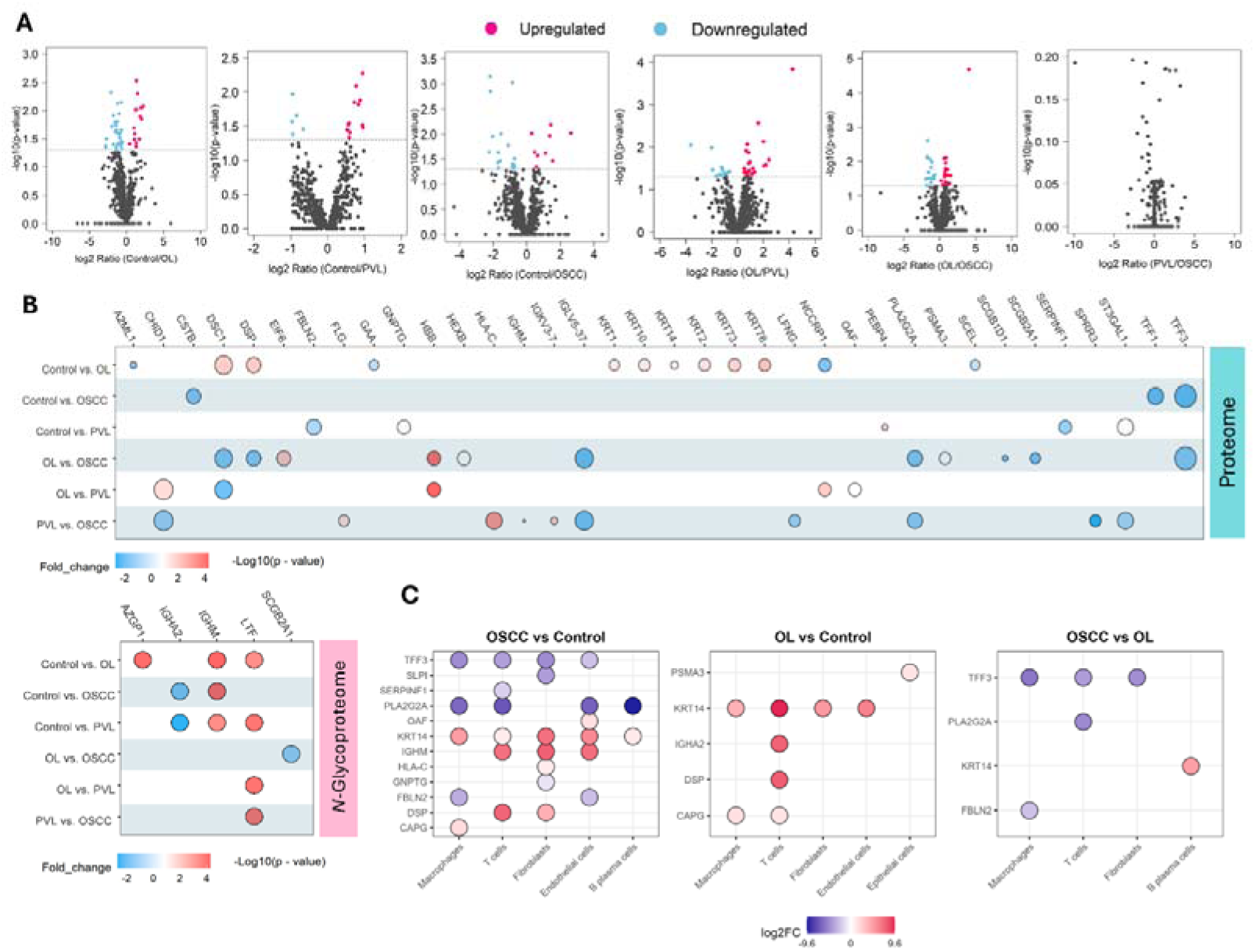
Tear fluid reveals dysregulated proteins across oral lesions. **(A)** Volcano plots of proteins differentially abundant in each pairwise comparison (Control vs OL, Control vs PVL, Control vs OSCC, OL vs PVL, OL vs OSCC, and PVL vs OSCC), tested with two-sided unpaired Student’s t-tests or Welch’s t-tests and corrected using the Benjamini–Hochberg test (P ≤ 0.05). Upregulated proteins are shown as red dots and downregulated proteins as blue dots. **(B)** Distribution of log2 fold changes for proteins and normalized intensity values of *N*-glycoproteins with differential abundance among groups. It was performed one-way ANOVA followed by Tukey’s post hoc test; P ≤ 0.05 for both datasets. Upregulated proteins are shown as red dots and downregulated proteins as blue dots. **(C)** Single-cell expression from the tissue of each differentially abundant protein identified in tear fluid across diferente cell types. Genes were considered differentially expressed when the adjusted P < 0.05 and the absolute log2 fold change > 1.

We next performed clustering analysis of the proteins with differential abundance from the tear fluid. We did not perform this analysis on *N*-glycoproteomic data because only five N-glycoproteins were differentially abundant across the groups. Clusters could not distinguish the clinical groups according to proteomic profiling (Fig. 5A). However, similar to the total proteins of the global proteome, the protein cluster was segregated into two different protein groups, for which we performed GO biological process analyses. Most of these proteins were enriched for epithelial cell differentiation (fig. S4). Notably, these proteins separated patients into three distinct clusters, indicating significant differences in the clinical and pathological features across them. For example, patient clusters were associated with oral epithelial dysplasia, immune infiltrate, and the presence of lymph-node metastasis (Fig. 5B). Epithelial dysplasia is the main histological predictor of malignant transformation (8), whereas lymph-node metastasis is an important component of staging that guides treatment and prognosis of OSCC (33). Other prognostic features also differed significantly across these clusters, such as nuclear pleomorphism, drop-shaped rete ridges, increased nuclear-cytoplasmic ratio, abnormal variation in nuclear size, and peritumoral desmoplasia (fig. S5). These results demonstrate that proteins with differential abundance were more closely associated with a range of clinicopathological features, reflecting disease biology.

**Figure 5.**
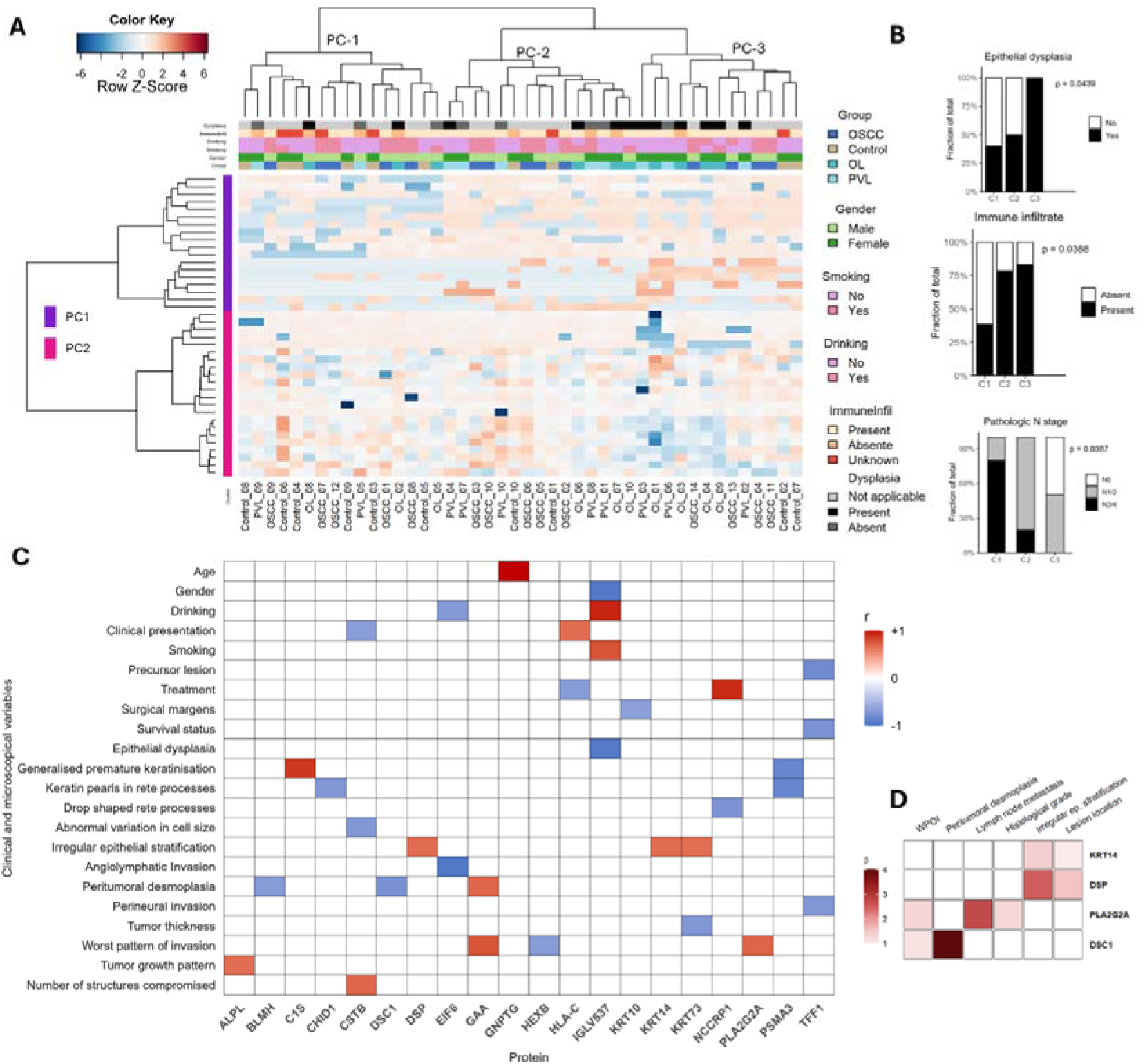
Differentially abundant proteins are associated with malignant transformation and prognosis of oral cancer. **(A)** LFQ intensity values of differentially abundant proteins were used for unsupervised hierarchical clustering analysis employing the *heatmap.3* function in R with Euclidean distance and Ward linkage. **(B)** Association between clinicopathological features and patient clustering (two-sided Fisher’s exact test; *P* ≤ 0.05). Bars represent the fraction (%) of patients by cluster. **(C)** Pearson’s *r* or Spearman’s ρ correlation analysis between LFQ intensity values and clinicopathological variables. Only significant correlations (*P* ≤ 0.05) with |r| or |ρ| ≥ ∼0.7 are shown. Red indicates positive correlation, and blue indicates negative correlation. The color scale reflects the correlation coefficient (–1 to +1). **(D)** Multiple linear regression indicating which clinicopathological variables were significant independent predictors of protein abundance (LFQ intensity values).

Considering that proteins with differential abundance were more closely associated with a range of clinicopathological features according to clustering analysis, we next sought to correlate these proteins with clinicopathological features using two distinct approaches. First, we measured the direction and strength of the relationship between individual protein abundance (LFQ intensity values) and clinicopathological data using Spearman’s ρ or Pearson’s *r* correlation. Second, we tested six multiple linear regression models to assess how many, and which, clinicopathological features could simultaneously predict protein abundance. These models were built using proteins (i) identified as statistically significant in the correlation analysis and (ii) with valid LFQ intensity values across all patients. Spearman’s ρ or Pearson’s correlation identified 20 proteins that were significantly associated with clinicopathological features (Fig. 5C). Correlations were considered significant when *P* ≤ 0.05 and |ρ| or |r| ≥ ∼0.7. These associations were related not only to malignant transformation but also to prognosis. Although several of these proteins have previously been linked to oral lesions at the tissue level, some have never been investigated in body fluids.

Four models reached statistical significance in the multiple linear regression analysis, demonstrating that protein abundance is not influenced by a single clinical or pathological feature, but by more than one simultaneously (table S8-11). Desmoplakin (DSP) and keratin 14 (KRT14) were influenced by irregular epithelial stratification and lesion location. In addition, the abundance of desmocollin 1 (DSC1) was influenced by the pattern of invasion and peritumoral desmoplasia, while the abundance of phospholipase A2 group IIA (PLA2G2A) was influenced by the presence of lymph-node metastasis, histological grade, and pattern of invasion (Fig. 5D). Although some of these proteins can be found in the corneal epithelium and have been previously detected in tear fluid (34), they are also present in the oral mucosa (35). KRT14, for instance, is present across all epithelial surfaces of the oral cavity, although it is more abundant in certain sites (35). This is consistent with our results showing that KRT14 abundance is influenced by epithelial stratification and lesion location.

Taking together, our results demonstrate that tear fluid can reflect the biology of oral diseases, opening opportunities for its use as a source of biomarkers.

## DISCUSSION

Cancer is a systemic disease that causes alterations in the structure and composition of tissues and body fluids (36,37). Cells from the tumor and tissue microenvironment releases substances into the systemic circulation that can be detected in sites distant from the tumor (36,37). These molecules can aid in screening, early diagnosis, prognosis and monitoring of cancer (36,37). In the early stages of carcinogenesis, tissue alterations are minimal and cannot be detected by routine histological assessment. It remains unclear whether such changes can be identified in body fluids it is not clear yet. Here, mass spectrometry-based discovery proteomics was employed to uncover alterations in the tear fluid of patients with oral lesions that have potential to become malignant. We demonstrated that dysregulation in the tear fluid proteome reflects patient features associated with malignant transformation and oral cancer prognosis.

The global proteome of tear fluid retrieved proteins associated with immune response. Most proteins, especially *N*-glycoproteins, were enriched for biological processes related to immune responses (Fig. 2C). In fact, tear fluid is essential for protecting the ocular surface against microorganisms (17,19). However, the overrepresentation of immune-enriched events may also reflect a systemic and coordinated response to early tumorigenesis during the immunosurveillance. Tissue cells can recruit peripheral immune cells to counteract tumor initiation and the secreted content from both tumor and immune cells can be detected in distant sites (36,37). Interestingly, a recent study identified that proteins relevant to blood and immune cells found in tear fluid are more likely to be transported by small EVs, and these EVs have a close connection with systemic circulation (21). This suggests that the proteome of tear fluid may reflect local and systemic immune response.

Besides, global proteome indicated differences in histological features from the patients based on clustering analysis. Unsurprisingly, grouping analysis did not segregate patients according to their clinical conditions. This suggests a high inter- and intra-patient heterogeneity also observed in other body fluids, such as saliva (38, 39). However, clustering analysis identified differences in histological features among patients. For example, the presence of high mitotic activity in the epithelium and abnormal variation in cell size differed significantly across patients assigned to each cluster (Fig. 2D). Both histological features are associated with an increased risk of malignant transformation (28,29). Abnormal variation in cell size in OL is associated with genomic instability, which is one of the hallmarks of cancer (29, 40).

Tear fluid is highly enriched with proteins that orchestrate the immune system. Our deconvoluted proteome showed that B cell–related proteins were highly abundant in OL, PVL, and OSCC (Fig. 3A). Interestingly, the immune cell composition of healthy control samples was distinct from that of the lesions based on clustering analysis (Fig 3A). This suggests that the tear fluid proteome of these lesions shares similarities in immune profiling. According to our deconvolution analysis, we observed that elevated proportions of macrophages in tear fluid were correlated with increased angiolymphatic invasion (table S5). It is increasingly clear that macrophage-driven angiolymphatic invasion is essential for the establishment of pre-metastatic niches (42,43). These cells employ distinct mechanisms to attach to endothelial cells, thereby promoting vascular invasion and tumor cell dissemination (41, 42). Notably, 12 out of 14 OSCC patients in our study presented lymph-node metastasis (table S1). Together, these results suggest that the global proteome of tear fluid reflects the stepwise progression toward malignancy and the maintenance of cancer in more advanced stages.

We identified 40 proteins that were differentially abundant across clinical conditions and could play a role in the biology of malignant transformation (table S7). It is beyond our aim to explore the role of each one, but it is worth noting they have been previously reported in tissues as drivers of malignancy and tumor progression. For example, two desmosomal proteins DSC1 and DSP, were dysregulated in tear fluid. These proteins are essential for maintaining cell adhesion, preventing invasion of epithelial cells into connective tissue, and metastasis. Both have been associated with the malignant progression of oral lesions to cancer, but not in biofluids (43, 44). We observe that these proteins are more abundant in healthy controls than OL, in line with their function of preventing cell decohesion. Conversely, they are found more abundant in OSCC compared to OL (Fig. 4B). These differences may be attributed to varying levels of protein abundance or expression across different stages of the disease.

In advanced tumors, cell adhesion proteins may be more highly expressed as tumor cells establish metastatic niches, thereby favoring dissemination. For instance, in colorectal cancer, strong tissue expression of DSC1 has been observed in budding cells at the tumor invasive front (45). Interestingly, our multivariate regression analysis revealed that DSC1 abundance was significantly higher in OSCC samples with tumor satellites located more than 1 mm away from the main tumor mass compared with samples displaying less invasive patterns (WPOI-V) (Fig. 5D and table S10). Our results suggest that alterations in cell adhesion that represent early and advanced stages of diseases could release proteins in body fluids favoring their detection in distant sites from the lesions. Therefore, tear fluid represents a promising liquid biopsy for tracking both early and advanced stages of oral malignant transformation.

Although this study has limitations regarding the sample size due to strict criteria to select the patients, such as excluding patients with systemic or ocular diseases, previous history of cancer, ongoing/previous history of cancer treatment and/or age under 40 years, it reveals that potential drivers of malignant progression are also present in tear fluid. For example, strong tissue expression of desmocollin 1 (DSC1) has been considered a predictor of epithelial dysplasia in oral lichen planus, a lesion that can also progress toward malignant transformation (43). In our data, DSC1 was upregulated in PVL and OSCC compared to OL. Of note, PVL is more prone to malignant progression than OL. Desmosomal proteins, such as desmoplakin (DSP), have been investigated in carcinogenesis because loss of cell adhesion is essential for tumor development and spreading (44). The tissue expression of DSP increases in the initial stages of OL but decreases in advanced stages of disease progression and in metastatic lymph nodes (44). In line with these results, the abundance of DSP was significantly decreased in OL compared to healthy controls in our data.

Altogether, this is the first study to characterize the tear-fluid proteome of patients with oral lesions, opening avenues to explore this easily accessible biofluid as a promising source of biomarkers for oral diseases. We performed a comprehensive statistical analysis to correlate tear fluid protein abundance with the clinical and pathological features of patients, providing robust evidence for these associations and capturing intra- and inter-patient variability as well as disease heterogeneity.

## MATERIALS AND METHODS

### Cohort and collection of tear fluid samples

This study was conducted in accordance with approved guidelines and the Declaration of Helsinki, and written informed consent was obtained from each patient. The Ethics Review Boards of both institutions—the Instituto do Câncer de São Paulo, Faculty of Medicine, University of São Paulo (ICESP/FM-USP), and the Piracicaba Dental School, University of Campinas (FOP/UNICAMP)—approved the study via Plataforma Brasil under protocol numbers CAAE 61402116.8.0000.0065 and CAAE 27338619.0.0000.5418, respectively. Tear fluid was collected from 44 patients: OL (n = 10), PVL (n = 10), OSCC (n = 14), and healthy controls (n = 10). The exclusion criteria were as follows: (a) ongoing treatment for OSCC; (b) lesions outside the oral cavity; and (c) contagious infectious diseases. Patients with a history of ocular disorders (e.g., Sjögren’s syndrome, dry eye syndrome, diabetic retinopathy) or other conditions (ocular surgery, trauma, contact lens use) were excluded. Patients were informed about the procedure and instructed to avoid blinking and physical stress, which could alter the chemical composition of tear fluid (46). We instilled 60 μL of 0.9% sodium chloride (saline) onto the ocular surface without direct contact to avoid irritation. Glass microcapillaries were used to collect tear fluid from each eye. Samples were stored at – 80 °C until analysis.

### Sample preparation for LC-MS/MS analysis

Randomization was performed in R (version 4.2.2) to avoid bias during sample preparation. Protein concentrations were measured using the Bradford assay (Bio-Rad, Hercules, CA, USA). After measurement, the total volume of each sample was adjusted to the same volume. Protein (10 µg) was denatured with 8 M urea (1:1), reduced with 5 mM DTT, alkylated with 14 mM iodoacetamide (IAA), and then treated with 5 mM DTT for 15 min at room temperature to quench the IAA. Urea was diluted to a final concentration of 1.6 M, followed by the addition of 1 mM calcium chloride. Proteins were digested with 0.2 µg of trypsin for 16 h at 37 °C; then an additional 0.2 µg of trypsin was added, and digestion continued for another 6 h at 37 °C. The reaction was quenched with 0.4% trifluoroacetic acid, and peptides were desalted using C18 StageTips (3M, USA) (47), dried in a vacuum concentrator, and stored at –20 °C.

### Mass spectrometry analysis using data-dependent acquisition

Forty-four LC-MS/MS runs were performed in total (one per patient) (table S12). Peptides were resuspended in 10 µL of 0.1% formic acid, and 2.0 µg of the peptide mixture was analyzed on an Orbitrap Velos 480 (Thermo Electron Scientific) coupled to a Protana nanospray ion source and a self-packed 8 cm × 75 µm i.d. Phenomenex Jupiter 10 µm C18 reversed-phase capillary column. Peptides were separated over a 60 min gradient from 2% to 90% acetonitrile in 0.1% formic acid at 300 nL/min. The nanoelectrospray voltage was 2.2 kV, and the source temperature was 275 °C. All methods were configured for data-dependent acquisition in positive-ion mode. Full-scan MS spectra (m/z 300–1600) were acquired in the Orbitrap analyzer after ion accumulation to a target value of 1 × 10^6 at a resolution of 60,000. The 20 most intense peptide ions with charge states ≥ 2 were sequentially isolated in a 3 m/z window to a target value of 5,000 and fragmented in the linear ion trap by low-energy collision-induced dissociation (normalized collision energy 35%). The signal threshold for triggering an MS/MS event was 1,000 counts. Dynamic exclusion was enabled with a list size of 500, an exclusion duration of 60 s, and a repeat count of 1. An activation Q of 0.25 and an activation time of 10 ms were used.

### Quality control of proteomics analysis

Quality evaluation of the LC–MS/MS runs was performed by measuring deviations in the retention times of three trypsin fragment peaks (m/z 421.7584, 2+; m/z 523.2855, 2+; and m/z 1106.0557, 2+). In addition, three iRT peptides were spiked into tear samples before injection to verify retention time and intensity, with intensities normalized by mean per sample and by mean per group. The iRT peptides were SSAAPPPPPR (iRT_Peptide_1), GISNEGQNASIK (iRT_Peptide_2), and HVLTSIGEK (iRT_Peptide_3) (Pierce™ Peptide Retention Time Calibration Mixture, Thermo Scientific, USA). Data were analyzed using Skyline 19.1 (48).

### Proteomic data analysis

We used a precursor mass tolerance of 4.5/6.0/10 ppm and a fragment-ion tolerance of 0.5 Da. Carbamidomethylation of cysteine was set as a fixed modification, with oxidation of methionine and N-terminal acetylation as variable modifications, and enzyme specificity set to Trypsin/P allowing up to two missed cleavages. A 1 % FDR was enforced at both the peptide and protein levels using a reverse-sequence decoy strategy. Protein grouping was performed by the Andromeda engine using the parsimony principle. Protein quantification was performed using the LFQ algorithm implemented in MaxQuant with razor + unique peptides. A minimal ratio counts of one and a 2-minute window for matching between runs were required for quantification. Proteins identified as “Reverse sequences” or “Only identified by site” were excluded from downstream analysis. Because keratins are intrinsic to the biology of OL, PVL, and OSCC, keratin peptides/proteins were retained and not filtered as “contaminants” (49,50); other common proteins listed as contaminants were removed. LFQ intensity values, which are normalized spectral intensities, were log□-transformed, and the dataset was then filtered to require a minimum of three valid values in at least one group. Missing LFQ values were imputed (zero replacement) for the proteome and N-glycoproteome hierarchical clustering and heatmap analyses. Perseus v2.1.3.0 was used to identify proteins differentially abundant across clinical groups (one-way ANOVA; Tukey’s post hoc test; P < 0.05). It was also used for two-sample comparisons: Control vs. OL; Control vs. PVL; and OL vs. PVL (two-sided unpaired Student’s t-test followed by Benjamini–Hochberg correction; P < 0.05), and Control vs. OSCC; OL vs. OSCC; and PVL vs. OSCC (two-sided unpaired Welch’s t-test followed by Benjamini–Hochberg correction; P < 0.05).

### *N*-glycopeptide search

The LC–MS/MS data were searched with the HCP workflow using Byonic v2.6.46 (Byos Software, Protein Metrics Inc., CA, USA) (23), with precursor and product ion mass tolerances of 10 ppm and 20 ppm, respectively. Trypsin-specific cleavages were applied, allowing up to two missed cleavages per peptide. Cysteine carbamidomethylation (+57.021 Da) was set as a fixed modification, with the following variable modifications: methionine oxidation (+15.994 Da) and N-glycosylation of sequon-localized Asn using compositions from the mammalian N-glycan database. A maximum of two common modifications and one rare modification were permitted. Data were searched against a UniProtKB database of human proteins (96 510 sequences; released June 25, 2021). All identifications were filtered to < 1% false-discovery rate (FDR) at the protein level and 0% at the peptide level, using a reverse-sequence decoy database (51). Only N-glycopeptides with PEP 2D scores < 0.001 were retained (52); low-confidence glycopeptides and those matched to reverse sequences were excluded. Glycopeptides were then manually grouped by summing the intensities of glycopeptide spectral matches (glycoPSMs) sharing the same UniProtKB ID, glycosylation site, and glycan composition. Unique glycopeptide abundances were normalized by dividing each glycopeptide’s intensity by the total intensity sum of its sample, correcting for inter-sample variation.

### Flow cytometry analysis

Tear fluid samples (∼80 µL) from 32 patients were collected in sterile physiological saline solution (0.9% NaCl) and maintained on ice until processing. The cells were washed with 500 µL of phosphate-buffered saline (PBS 1×) by centrifugation at 400 × g for 5 min at room temperature (RT). For Fc receptor blocking, 0.5 µL of Fc Block reagent was added to 9.5 µL of PBS 1× per sample, and the cells were incubated for 10 min at RT. Surface staining was then performed using 1 µL of PE-Cy7–conjugated anti-human CD19 monoclonal antibody (clone HIB19, BD Pharmingen™, Cat. No. 560728) diluted in 50 µL of PBS 1×. The cells were incubated for 30 min at RT (protected from light). After staining, the cells were washed with 500 µL of PBS 1× (400 × g, 5 min, RT). The cells were then fixed with 1 mL of 4% paraformaldehyde (PFA) for 10 min at RT, followed by centrifugation at 400 × g for 5 min. Subsequently, the cells were washed again with 500 µL of PBS 1× (400 × g, 5 min, RT) and finally resuspended in approximately 200 µL of filtered PBS 1× for acquisition. Flow-cytometric acquisition was performed on a BD FACSCanto II cytometer (BD Biosciences, USA). At least 30,000 gated events were acquired assuring the reliability of positive populations. The cells were then gated on SSC-A x FSC-A to exclude debris and FSC-H x FSC-A to select single cells. Data acquisition and fluorescence intensity analysis were carried out using FlowJo v10.8.1 software (BD Biosciences). Statistical analyses were performed in GraphPad Prism v8.2.1 (GraphPad Software, USA). Differences among experimental groups were evaluated using ordinary one-way ANOVA, followed by Tukey’s multiple comparisons test. Statistical significance was defined as P ≤ 0.05.

### Single-cell analysis of tissue samples using public dataset

The scRNA-seq processed data were obtained and further preprocessed as described in the literature and by the dataset authors (31, 53). The original dataset consisted of 23 patients with head and neck cancer; however, after excluding OSCC HPV+ samples as well as samples from non-oral cavity sites, we analyzed data from 17 patients. The integration step, as well as the identification and removal of doublets, had already been performed in the processed data available (31). Filters were applied to remove genes expressed in less than 0.1% of cells. Moreover, cells with >200 genes, <8,000 genes, and <10% of mitochondrial gene expression in unique molecular identifier (UMI) counts were retained (31). An additional filter was introduced to remove cells with less than 5% ribosomal reads (54). In addition, MALAT1, mitochondrial, and hemoglobin genes were excluded (55). All filtering procedures mentioned above were performed using the following packages: numpy v1.23.5, pandas v1.4.4, and scanpy v1.7.2. Each cell was normalized to 10,000 UMI and log□-transformed. To reduce dimensionality, the *highly_variable_genes* function was used to calculate the 11,569 (half of the total number of cells) highly variable genes selected for downstream analysis (54). Next, we performed differential expression analysis using *scvi-tools* v0.14.5 powered by PyTorch v2.0.0, and the *rank_genes_groups* function from Scanpy. In both methods, all pairwise comparisons were tested among the following conditions: normal tissue (NL), primary cancer (CA), and precancerous leukoplakia (LP). Genes were considered differentially expressed when the adjusted *P*-value was <0.05 and the absolute log(fold change) >1. All scRNA-seq analyses described in this work were performed using Python v3.8.16, and all visualizations were created with the Matplotlib package v3.6.3. The analyses were conducted on a workstation with the following configuration: Intel Core i7-4790K 4 GHz Quad-Core CPU, 32 GB DDR3-1600 MHz RAM, and GeForce RTX 3060 12 GB GPU.

### GO annotation of biological processes

Gene Ontology (GO) biological processes significantly enriched in each cluster of the global proteome, N-glycoproteome, and differential proteome were evaluated using the Pan-GO Functionome, the most recent and expanded version of the PANTHER database (56). Protein and N-glycoprotein lists (gene names) from each cluster were uploaded to the Pan-GO Functionome for mapping and enrichment of GO biological processes. Statistical enrichment was assessed using Fisher’s exact test followed by Benjamini–Hochberg correction for multiple hypotheses (*adjusted P* ≤ 0.05). GO annotations were based on the database version released in August 2025.

### Association of patient clustering analysis with clinicopathological data

The association between patient clusters and clinicopathological data was evaluated in RStudio using the stats package (v4.4.3). Two-sided Fisher’s exact tests (P ≤ 0.05) were applied to assess these associations.

### Correlation of proteomic data with clinicopathological features

Pearson’s *r* or Spearman’s ρ correlation analyses were performed for all proteins showing differential abundance to evaluate their association with clinicopathological variables. LFQ intensity values (dependent variables) were correlated with clinicopathological features (independent variables) using R software (v4.0.4). Correlations were considered significant when *P* ≤ 0.05 and |*r*| or |ρ| ≥ ∼0.7. The clinicopathological features analyzed are described in detail in table S5.

### Multiple linear regression analysis

We performed multiple linear regression with backward elimination to assess the relationship between the dependent variable (LFQ intensity values) and a set of independent variables (clinicopathological data). Our objective was to predict how clinical and pathological features collectively influence protein abundance. An individual model was built for each protein, considering the following inclusion criteria: (1) proteins with complete LFQ intensity values across all patients in each clinical group; and (2) proteins that showed significant correlations in the Pearson/Spearman correlation analysis. The clinical and pathological features included in the models are detailed in Table S5. Initially, a general model was constructed using features common to all clinical groups, including mean age (≥ vs. <45 years; ≥ vs. <60 years), gender (male/female), smoking habits (yes/no), alcohol consumption (yes/no), lesion location (tongue/other), and immune cell infiltrate based on histological evaluation (present/absent). Subsequent models were adjusted to include features specific to each clinical group. For example, cytological and architectural features of epithelial dysplasia were included only for the OL/PVL group, whereas peritumoral desmoplasia, lymph-node metastasis, and perineural invasion were included exclusively for OSCC patients. Model diagnostics included assessment of residuals, normality, multicollinearity, and homoscedasticity. Multicollinearity among independent variables was considered acceptable when tolerance values were close to 1 and variance inflation factor (VIF) values were <5. Autocorrelation was evaluated using the Durbin–Watson test (*p* ≤ 0.05).

### Statistical analysis

Statistical analyses were performed using R version 4.5.1, Jamovi 2.6.22.0 (https://www.jamovi.org), and GraphPad Prism version 8.0.1 (https://www.graphpad.com). The tests used in each analysis are indicated in the figure legends, tables, and the Materials and Methods section. One-way ANOVA followed by Tukey’s post hoc test was used for the differential abundance analysis of the proteome, N-glycoproteome, and flow cytometry analysis. Two-sided unpaired Student’s *t*-test or Welch’s *t*-test, followed by Benjamini–Hochberg correction, was used in pairwise comparisons in differential protein abundance analysis. Two-sided Fisher’s exact test was used for the association between clustering and clinicopathological data. Pearson and Spearman correlation, as well as multiple linear regression analysis, are described in detail in the Materials and Methods section. Statistical significance for all tests was established at *P* ≤ 0.05.

### Data availability

The mass spectrometry proteomics data generated in this study are available via the ProteomeXchange Consortium (56). The discovery dataset can be accessed under accession PXD042738. Public single-cell RNA-seq data used for computational deconvolution and tissue scRNA-seq were downloaded and analyzed from the Gene Expression Omnibus under accession GSE181919. In total, we downloaded 17 datasets: 3 normal tissues (GSM5514381, GSM5514375, GSM5514376), 4 oral leukoplakia (OL) (GSM5514384, GSM5514385, GSM5514386, GSM5514387), and 10 OSCC (primary cancer) (GSM5514352, GSM5514364, GSM5514366, GSM5514353, GSM5514354, GSM5514356, GSM5514357, GSM5514358, GSM5514361, GSM5514388).

## Supporting information

Previous version of the manuscript

## REFERENCES

1. Bittar RF, Ferraro HP, Ribas MH, Lehn CN. Predictive factors of occult neck metastasis in patients with oral squamous cell carcinoma. Braz J Otorhinolaryngol. 2016 Sep-Oct;82(5):543-7. doi: 10.1016/j.bjorl.2015.09.005. Epub 2015 Dec 17. PMID: 26749457; PMCID: PMC9444684.

2. Wang Y, Yang T, Gan C, Wang K, Sun B, Wang M, Zhu F. Temporal and spatial patterns of recurrence in oral squamous cell carcinoma, a single-center retrospective cohort study in China. BMC Oral Health. 2023 Sep 19;23(1):679. doi: 10.1186/s12903-023-03204-7. PMID: 37726764; PMCID: PMC10510235.

3. Henry M, Rosberger Z, Bertrand L, Klassen C, Hier M, Zeitouni A, Kost K, Mlynarek A, Richardson K, Black M, MacDonald C, Zhang X, Chartier G, Frenkiel S. Prevalence and Risk Factors of Suicidal Ideation among Patients with Head and Neck Cancer: Longitudinal Study. Otolaryngol Head Neck Surg. 2018 Nov;159(5):843–852. doi: 10.1177/0194599818776873. Epub 2018 Jun 5. PMID: 29865939.

4. Piemonte ED, Gilligan GM, Garola F, Lazos JP, Panico RL, Normando AGC, Santos-Silva AR, Warnakulasuriya S. Differences among oral carcinomas arising de novo from those associated with oral potentially malignant disorders: a systematic review and meta-analysis. Oral Surg Oral Med Oral Pathol Oral Radiol. 2024 Jun;137(6):613–631. doi: 10.1016/j.oooo.2024.03.006. Epub 2024 Mar 19. PMID: 38609795.

5. Iocca O, Sollecito TP, Alawi F, Weinstein GS, Newman JG, De Virgilio A, et al. Potentially malignant disorders of the oral cavity and oral dysplasia: A systematic review and meta-analysis of malignant transformation rate by subtype. Head Neck. 2020 Mar;42(3):539–555.

6. Hvid-Jensen F, Pedersen L, Drewes AM, Sørensen HT, Funch-Jensen P. Incidence of adenocarcinoma among patients with Barrett’s esophagus. N Engl J Med. 2011 Oct 13;365(15):1375–83. doi: 10.1056/NEJMoa1103042. PMID: 21995385.

7. Lodi G, Franchini R, Warnakulasuriya S, Varoni EM, Sardella A, Kerr AR, Carrassi A, MacDonald LC, Worthington HV. Interventions for treating oral leukoplakia to prevent oral cancer. Cochrane Database Syst Rev. 2016 Jul 29;7(7):CD001829. doi: 10.1002/14651858.CD001829.pub4. PMID: 27471845; PMCID: PMC6457856.

8. Odell E, Kujan O, Warnakulasuriya S, Sloan P. Oral epithelial dysplasia: Recognition, grading and clinical significance. Oral Dis. 2021 Nov;27(8):1947–1976. doi: 10.1111/odi.13993. Epub 2021 Sep 14. PMID: 34418233.

9. Thompson LDR, Fitzpatrick SG, Müller S, Eisenberg E, Upadhyaya JD, Lingen MW, Vigneswaran N, Woo SB, Bhattacharyya I, Bilodeau EA, Carlos R, Islam MN, Leon ME, Lewis JS Jr, Magliocca KR, Mani H, Mehrad M, Purgina B, Richardson M, Wenig BM, Cohen DM. Proliferative Verrucous Leukoplakia: An Expert Consensus Guideline for Standardized Assessment and Reporting. Head Neck Pathol. 2021 Jun;15(2):572–587. doi: 10.1007/s12105-020-01262-9. Epub 2021 Jan 7. PMID: 33415517; PMCID: PMC8134585.

10. Proaño-Haro A, Bagan L, Bagan JV. Recurrences following treatment of proliferative verrucous leukoplakia: A systematic review and meta-analysis. J Oral Pathol Med. 2021 Sep;50(8):820–828. doi: 10.1111/jop.13178. Epub 2021 Apr 13. PMID: 33765364.

11. Nakayasu ES, Gritsenko M, Piehowski PD, Gao Y, Orton DJ, Schepmoes AA, Fillmore TL, Frohnert BI, Rewers M, Krischer JP, Ansong C, Suchy-Dicey AM, Evans-Molina C, Qian WJ, Webb-Robertson BM, Metz TO. Tutorial: best practices and considerations for mass-spectrometry-based protein biomarker discovery and validation. Nat Protoc. 2021 Aug;16(8):3737–3760. doi: 10.1038/s41596-021-00566-6. Epub 2021 Jul 9. PMID: 34244696; PMCID: PMC8830262.

12. Sá JO, Trino LD, Oliveira AK, Lopes AFB, Granato DC, Normando AGC, Santos ES, Neves LX, Carnielli CM, Paes Leme AF. Proteomic approaches to assist in diagnosis and prognosis of oral cancer. Expert Rev Proteomics. 2021 Apr;18(4):261–284. doi: 10.1080/14789450.2021.1924685. Epub 2021 May 18. PMID: 33945368.

13. Surdu A, Foia LG, Luchian I, Trifan D, Tatarciuc MS, Scutariu MM, Ciupilan C, Budala DG. Saliva as a Diagnostic Tool for Systemic Diseases-A Narrative Review. Medicina (Kaunas). 2025 Jan 30;61(2):243. doi: 10.3390/medicina61020243. PMID: 40005360; PMCID: PMC11857487.

14. Tasoulas J, Patsouris E, Giaginis C, Theocharis S. Salivaomics for oral diseases biomarkers detection. Expert Rev Mol Diagn. 2016;16(3):285–95. doi: 10.1586/14737159.2016.1133296. Epub 2016 Jan 11. PMID: 26680995.

15. Nagler RM, Hershkovich O. Age-related changes in unstimulated salivary function and composition and its relations to medications and oral sensorial complaints. Aging Clin Exp Res. 2005 Oct;17(5):358–66. doi: 10.1007/BF03324623. PMID: 16392409.

16. Nicholson K, Liu W, Fitzpatrick D, Hardacre KA, Roberts S, Salerno J, Stranges S, Fortin M, Mangin D. Prevalence of multimorbidity and polypharmacy among adults and older adults: a systematic review. Lancet Healthy Longev. 2024 Apr;5(4):e287–e296. doi: 10.1016/S2666-7568(24)00007-2. Epub 2024 Mar 4. PMID: 38452787.

17. Adigal, S. S., Rizvi, A., Rayaroth, N. V., John, R. V., Barik, A., Bhandari, S., George, S. D., Lukose, J., Kartha, V. B., & Chidangil, S. (2021). Human tear fluid analysis for clinical applications: progress and prospects. Expert review of molecular diagnostics, 21(8), 767–787.

18. D, Y., Byrum, S. D., Acott, A. A., Siegel, E. R., Washam, C. L., Klimberg, V. S., & Mancino, A. T. (2022). Proteomic profiling of tear fluid as a promising non-invasive screening test for colon cancer. American journal of surgery, 224(1 Pt A), 19–24. 10.1016/j.amjsurg.2022.03.029

19. Gijs M, van de Sande N, Bonnet C, Schmeetz J, Fernandes R, Travé-Huarte S, Huertas-Bello M, Bo Chiang JC, Boychev N, Sharma S; Tear Research Network Scoping Review taskforce. A comprehensive scoping review of methodological approaches and clinical applications of tear fluid biomarkers. Prog Retin Eye Res. 2025 May;106:101338. doi: 10.1016/j.preteyeres.2025.101338. Epub 2025 Feb 13. PMID: 39954936.

20. Fatima A, Sanyal S, Jha GK, Kaliki S, Pallavi R. The enigmatic world of tear extracellular vesicles (EVs)-exploring their role in ocular health and beyond. FEBS Lett. 2025 May;599(10):1346–1372. doi: 10.1002/1873-3468.70004. Epub 2025 Feb 17. PMID: 39961136.

21. Hu L, Liu X, Zheng Q, Chen W, Xu H, Li H, Luo J, Yang R, Mao X, Wang S, Chen T, Lee LP, Liu F. Interaction network of extracellular vesicles building universal analysis via eye tears: iNEBULA. Sci Adv. 2023 Mar 17;9(11):eadg1137. doi: 10.1126/sciadv.adg1137. Epub 2023 Mar 15. PMID: 36921051; PMCID: PMC10017052.

22. Runström G, Mann A, Tighe B. The fall and rise of tear albumin levels: a multifactorial phenomenon. Ocul Surf. 2013 Jul;11(3):165–80. doi: 10.1016/j.jtos.2013.03.001. Epub 2013 May 1. PMID: 23838018.

23. Bern M, Kil YJ, Becker C. Byonic: advanced peptide and protein identification software. Curr Protoc Bioinformatics. 2012 Dec;Chapter 13:13.20.1-13.20.14. doi: 10.1002/0471250953.bi1320s40. PMID: 23255153; PMCID: PMC3545648.

24. Escher C, Reiter L, MacLean B, Ossola R, Herzog F, Chilton J, MacCoss MJ, Rinner O. Using iRT, a normalized retention time for more targeted measurement of peptides. Proteomics. 2012 Apr;12(8):1111–21. doi: 10.1002/pmic.201100463. PMID: 22577012; PMCID: PMC3918884.

25. Nguyen-Khuong T, Everest-Dass AV, Kautto L, Zhao Z, Willcox MD, Packer NH. Glycomic characterization of basal tears and changes with diabetes and diabetic retinopathy. Glycobiology. 2015 Mar;25(3):269–83. doi: 10.1093/glycob/cwu108. Epub 2014 Oct 9. PMID: 25303961.

26. Ponzini E. Tear biomarkers. Adv Clin Chem. 2024;120:69-115. doi: 10.1016/bs.acc.2024.03.002. Epub 2024 Apr 11. PMID: 38762243.

27. Jones G, Lee TJ, Glass J, Rountree G, Ulrich L, Estes A, Sezer M, Zhi W, Sharma S, Sharma A. Comparison of Different Mass Spectrometry Workflows for the Proteomic Analysis of Tear Fluid. Int J Mol Sci. 2022 Feb 19;23(4):2307. doi: 10.3390/ijms23042307. PMID: 35216421; PMCID: PMC8875482.

28. Rodrigues AZ, Laureano NK, Maraschin BJ, da Silva AD, da Silva VP, Rados PV, Visioli F. Diagnostic Criteria for Oral Epithelial Dysplasia: Predicting Malignant Transformation. Head Neck Pathol. 2025 Feb 7;19(1):21. doi: 10.1007/s12105-025-01754-6. PMID: 39918668; PMCID: PMC11806176.

29. Cai X, Zhang J, Zhang H, Zhou X, Zhou Z, Jing F, Luo H, Li T. Architectural and cytological features of epithelial dysplasia associated with transformation risk. Oral Dis. 2024 Jul;30(5):3028–3038. doi: 10.1111/odi.14809. Epub 2023 Nov 20. PMID: 37983891.

30. Newman AM, Steen CB, Liu CL, Gentles AJ, Chaudhuri AA, Scherer F, Khodadoust MS, Esfahani MS, Luca BA, Steiner D, Diehn M, Alizadeh AA. Determining cell type abundance and expression from bulk tissues with digital cytometry. Nat Biotechnol. 2019 Jul;37(7):773–782. doi: 10.1038/s41587-019-0114-2. Epub 2019 May 6. PMID: 31061481; PMCID: PMC6610714.

31. Choi JH, Lee BS, Jang JY, Lee YS, Kim HJ, Roh J, Shin YS, Woo HG, Kim CH. Single-cell transcriptome profiling of the stepwise progression of head and neck cancer. Nat Commun. 2023 Feb 24;14(1):1055. doi: 10.1038/s41467-023-36691-x. PMID: 36828832; PMCID: PMC9958029.

32. Fu ZM, Bao YY, Dai LB, Zhong JT, Chen HC, Chen Z, Zhou SH. Comprehensive Single-Cell RNA Atlas of Human Laryngeal Normal, Preneoplastic, and Tumorigenic States. Clin Cancer Res. 2025 Aug 1;31(15):3332–3343. doi: 10.1158/1078-0432.CCR-24-3679. PMID: 40407728; PMCID: PMC12314508.

33. Almangush A, Mäkitie AA, Triantafyllou A, de Bree R, Strojan P, Rinaldo A, Hernandez-Prera JC, Suárez C, Kowalski LP, Ferlito A, Leivo I. Staging and grading of oral squamous cell carcinoma: An update. Oral Oncol. 2020 Aug;107:104799. doi: 10.1016/j.oraloncology.2020.104799. Epub 2020 May 20. PMID: 32446214.

34. Ahmed S, Altman J, Jones G, Lee TJ, Robertson DM, Zhi W, Sharma S, Sharma A. Mass spectrometric detection of keratins in tear fluid. Exp Eye Res. 2025 Feb;251:110231. doi: 10.1016/j.exer.2025.110231. Epub 2025 Jan 4. PMID: 39761842; PMCID: PMC11798696.

35. Su L, Morgan PR, Lane EB. Keratin 14 and 19 expression in normal, dysplastic and malignant oral epithelia. A study using in situ hybridization and immunohistochemistry. J Oral Pathol Med. 1996 Jul;25(6):293–301. doi: 10.1111/j.1600-0714.1996.tb00265.x. PMID: 8887072.

36. Crosby D, Bhatia S, Brindle KM, Coussens LM, Dive C, Emberton M, Esener S, Fitzgerald RC, Gambhir SS, Kuhn P, Rebbeck TR, Balasubramanian S. Early detection of cancer. Science. 2022 Mar 18;375(6586):eaay9040. doi: 10.1126/science.aay9040. Epub 2022 Mar 18. PMID: 35298272.

37. Hiam-Galvez KJ, Allen BM, Spitzer MH. Systemic immunity in cancer. Nat Rev Cancer. 2021 Jun;21(6):345–359. doi: 10.1038/s41568-021-00347-z. Epub 2021 Apr 9. PMID: 33837297; PMCID: PMC8034277.

38. Arroyo E, Pérez Sayáns M, Bravo SB, de Oliveira Barbeiro C, Paravani Palaçon M, Chamorro Petronacci CM, García Vence M, Chantada Vázquez MDP, Blanco Carrión A, Suárez Peñaranda JM, García García A, Gándara Vila P, Días Almeida J, Veríssimo da Costa GC, Sousa Nogueira FC, Medeiros Evaristo JA, de Abreu Pereira D, Rintala M, Salo T, Rautava J, Padín Iruegas E, Oliveira Alves MG, Morandin Ferrisse T, Albergoni da Silveira H, Esquiche León J, Vilela Silva E, Flores IL, Bufalino A. Identification of Proteomic Biomarkers in Proliferative Verrucous Leukoplakia through Liquid Chromatography With Tandem Mass Spectrometry. Lab Invest. 2023 Oct;103(10):100222. doi: 10.1016/j.labinv.2023.100222. Epub 2023 Jul 26. PMID: 37507024.

39. Sivadasan P, Gupta MK, Sathe G, Sudheendra HV, Sunny SP, Renu D, Hari PS, Gowda H, Suresh A, Kuriakose MA, Sirdeshmukh R. Salivary proteins from dysplastic leukoplakia and oral squamous cell carcinoma and their potential for early detection. J Proteomics. 2020 Feb 10;212:103574. doi: 10.1016/j.jprot.2019.103574. Epub 2019 Nov 7. PMID: 31706945.

40. Negrini S, Gorgoulis VG, Halazonetis TD. Genomic instability--an evolving hallmark of cancer. Nat Rev Mol Cell Biol. 2010 Mar;11(3):220–8. doi: 10.1038/nrm2858. PMID: 20177397.

41. Fu LQ, Du WL, Cai MH, Yao JY, Zhao YY, Mou XZ. The roles of tumor-associated macrophages in tumor angiogenesis and metastasis. Cell Immunol. 2020 Jul;353:104119. doi: 10.1016/j.cellimm.2020.104119. Epub 2020 May 4. PMID: 32446032.

42. Doak GR, Schwertfeger KL, Wood DK. Distant Relations: Macrophage Functions in the Metastatic Niche. Trends Cancer. 2018 Jun;4(6):445–459. doi: 10.1016/j.trecan.2018.03.011. Epub 2018 May 3. PMID: 29860988; PMCID: PMC5990045.

43. Mattila R, Alanen K, Syrjänen S. Desmocollin expression in oral atrophic lichen planus correlates with clinical behavior and DNA content. J Cutan Pathol. 2008 Sep;35(9):832–8. doi: 10.1111/j.1600-0560.2007.00903.x. Epub 2008 Apr 18. PMID: 18422976.

44. Sawant S, Dongre H, Ahire C, Sharma S, Jamghare S, Kansara Y, Rane P, Kanojia D, Patil A, Chaukar D, Gupta S, D’Cruz A, Vaidya M, Dongre P. Alterations in desmosomal adhesion at protein and ultrastructure levels during the sequential progressive grades of human oral tumorigenesis. Eur J Oral Sci. 2018 Aug;126(4):251–262. doi: 10.1111/eos.12426. Epub 2018 Jun 15. PMID: 29905981.

45. Bartolomé RA, Pintado-Berninches L, Martín-Regalado Á, Robles J, Calvo-López T, Ortega-Zapero M, Llorente-Sáez C, Boukich I, Fernandez-Aceñero MJ, Casal JI. A complex of cadherin 17 with desmocollin 1 and p120-catenin regulates colorectal cancer migration and invasion according to the cell phenotype. J Exp Clin Cancer Res. 2024 Jan 24;43(1):31. doi: 10.1186/s13046-024-02956-6. PMID: 38263178; PMCID: PMC10807196.

46. Dumortier, G., & Chaumeil, J. C. (2004). Lachrymal determinations: methods and updates on biopharmaceutical and clinical applications. Ophthalmic research, 36(4), 183–194. 10.1159/000078776.

47. Rappsilber J, Mann M, Ishihama Y. Protocol for micro-purification, enrichment, pre-fractionation and storage of peptides for proteomics using StageTips. Nat Protoc. 2007;2(8):1896–906. doi: 10.1038/nprot.2007.261. PMID: 17703201.

48. MacLean B, Tomazela DM, Shulman N, Chambers M, Finney GL, Frewen B, Kern R, Tabb DL, Liebler DC, MacCoss MJ. Skyline: an open source document editor for creating and analyzing targeted proteomics experiments. Bioinformatics. 2010 Apr 1;26(7):966–8. doi: 10.1093/bioinformatics/btq054. Epub 2010 Feb 9. PMID: 20147306; PMCID: PMC2844992.

49. Woods RSR, Callanan D, Jawad H, Molony P, Werner R, Heffron C, Feeley L, Sheahan P. Cytokeratin 7 and 19 expression in oropharyngeal and oral squamous cell carcinoma. Eur Arch Otorhinolaryngol. 2022 Mar;279(3):1435–1443. doi: 10.1007/s00405-021-06894-3. Epub 2021 May 27. PMID: 34046748.

50. Murthy O G, Lau J, Balasubramaniam R, Frydrych AM, Kujan O. Unraveling the Keratin Expression in Oral Leukoplakia: A Scoping Review. Int J Mol Sci. 2024 May 21;25(11):5597. doi: 10.3390/ijms25115597. PMID: 38891785; PMCID: PMC11172080.

51. Bern MW, Kil YJ. Two-dimensional target decoy strategy for shotgun proteomics. J Proteome Res. 2011 Dec 2;10(12):5296–301. doi: 10.1021/pr200780j. Epub 2011 Nov 7. PMID: 22010998; PMCID: PMC3230778.

52. Kawahara R, Recuero S, Srougi M, Leite KRM, Thaysen-Andersen M, Palmisano G. The Complexity and Dynamics of the Tissue Glycoproteome Associated With Prostate Cancer Progression. Mol Cell Proteomics. 2021;20:100026. doi: 10.1074/mcp.RA120.002320. Epub 2021 Jan 5. PMID: 33127837; PMCID: PMC8010466.

53. Melms, J.C., Biermann, J., Huang, H. et al. A molecular single-cell lung atlas of lethal COVID-19. Nature 595, 114–119 (2021). 10.1038/s41586-021-03569-1.

54. Dong B, Miao J, Wang Y, Luo W, Ji Z, Lai H, Zhang M, Cheng X, Wang J, Fang Y, Zhu HH, Chua CW, Fan L, Zhu Y, Pan J, Wang J, Xue W, Gao WQ. Single-cell analysis supports a luminal-neuroendocrine transdifferentiation in human prostate cancer. Commun Biol. 2020 Dec 16;3(1):778. doi: 10.1038/s42003-020-01476-1. PMID: 33328604; PMCID: PMC7745034.

55. Zhao C, Biondic S, Vandal K, Björklund ÅK, Hagemann-Jensen M, Sommer TM, Canizo J, Clark S, Raymond P, Zenklusen DR, Rivron N, Reik W, Petropoulos S. Single-cell multi-omics of human preimplantation embryos shows susceptibility to glucocorticoids. Genome Res. 2022 Sep 27;32(9):1627–1641. doi: 10.1101/gr.276665.122. PMID: 35948369; PMCID: PMC9528977.

56. Perez-Riverol Y, Csordas A, Bai J, et al. The PRIDE database and related tools and resources in 2019: improving support for quantification data. Nucleic Acids Res. 2019;47(D1):D442–D450. doi:10.1093/nar/gky1106

