## Supplementary material for "Tear fluid as noninvasive liquid biopsy reveals proteins associated with malignant transformation of oral lesions": Previous version of the manuscript

bioRxiv preprint doi: https://doi.org/10.1101/2025.11.06.686738; this version posted November 7, 2025. The copyright holder for this preprint (which was not certified by peer review) is the author/funder. All rights reserved. No reuse allowed without permission.

### Tear fluid as noninvasive liquid biopsy reveals proteins associated with malignant transformation of oral lesions

Erison S. Santos[1,2], Daniela C. Granato[1], Carolina M. Carnielli[1], Ariane F. B. Lopes[1], Daniella de Figueiredo[1], Luciana D. Trino[1], Fabio M. S Patroni[1], Bianca A. Pauletti[1], Romênia R. Domingues[1], Jamile Sa[1], Tatiane M. Rossi[1], Ana G.C. Normando[1,2], Narmeen Daher[3], Nataly Kravchenko-Balasha[3], Paul Debasis[4], Rosane Minghim[4], Luiz P. Kowalski [5,6], Alan R. Santos-Silva[2], Márcio A. Lopes[2], Thaís B. Brandão[7], Ana C. Prado-Ribeiro[7], Adriana F. Paes Leme[1]*

[1]Laboratório Nacional de Biociências - LNBio, Centro Nacional de Pesquisa em Energia e Materiais - CNPEM, Campinas, SP 13083-100, Brazil.

[2]Departamento de Diagnóstico Oral, Faculdade de Odontologia de Piracicaba, Universidade Estadual de Campinas - UNICAMP, Piracicaba, SP 13414-903, Brazil.

[3]Institute of Dental Sciences, Faculty of Dental Medicine, The Hebrew University of Jerusalem, P.O.B. 12272, Ein Kerem, Jerusalem, ISRAEL 91120

[4]School of Computer Science and Information Technology, University College Cork, Cork, Ireland

[5] Head and Neck Surgery Department and LIM 28, University of São Paulo Medical School (FMUSP), São Paulo, Brazil

[6] Department of Head and Neck Surgery and Otorhinolaryngology, A.C. Camargo Cancer Center, São Paulo, Brazil

[7]Instituto do Câncer do Estado de São Paulo - ICESP, Faculdade de Medicina, Universidade de São Paulo - USP, São Paulo, SP 01246-000, Brazil

Short title: Tear fluid reflects oral malignant transformation

Disciplines: Biochemistry; Oncology

*Corresponding author**:**

bioRxiv preprint doi: https://doi.org/10.1101/2025.11.06.686738; this version posted November 7, 2025. The copyright holder for this preprint (which was not certified by peer review) is the author/funder. All rights reserved. No reuse allowed without permission.


#### Abstract

Oral leukoplakias (OLs) are premalignant lesions that can progress into oral squamous cell carcinoma (OSCC). This study hypothesized that tear fluid, as a noninvasive biofluid, reflects proteomic alterations associated with malignant transformation. The tear proteome of 44 individuals, including healthy controls, OL/PVL (proliferative verrucous leukoplakia), and OSCC patients, was deeply profiled, revealing 828 protein groups clustered according to histopathological alterations. N-glycoproteome analysis identified immune-related proteins, while public RNA-seq integration indicated immune imbalance marked by increased B-cell and decreased macrophage signatures during disease progression. Several immune-associated proteins and epithelial markers, including desmoplakin, KRT14, and DSC1, emerged as potential indicators of malignant transformation. These findings demonstrate that tear fluid reflects oral carcinogenic processes, thereby serving as a noninvasive liquid biopsy for early detection and clinical monitoring.

bioRxiv preprint doi: https://doi.org/10.1101/2025.11.06.686738; this version posted November 7, 2025. The copyright holder for this preprint (which was not certified by peer review) is the author/funder. All rights reserved. No reuse allowed without permission.

Advances in mass spectrometry-based proteomics have driving the discovery of biomarkers through noninvasive liquid biopsy (11). In the oral cavity, saliva is recognized as a promising type of liquid biopsy due to its proximity to oral lesions,

bioRxiv preprint doi: https://doi.org/10.1101/2025.11.06.686738; this version posted November 7, 2025. The copyright holder for this preprint (which was not certified by peer review) is the author/funder. All rights reserved. No reuse allowed without permission.

Unstimulated tear fluid samples from 44 individuals were collected from patients with oral leukoplakia (OL, N=10), with proliferative verrucous leukoplakia (PVL, N=10), with oral cancer (OSCC, N= 14) and from healthy controls (N=10) (table S1), and the global proteome and its respective non-enriched *N*-glycoproteome were obtained (Fig. 1A) (23). The statistically significant proteins and *N*-glycopeptides were associated with

bioRxiv preprint doi: https://doi.org/10.1101/2025.11.06.686738; this version posted November 7, 2025. The copyright holder for this preprint (which was not certified by peer review) is the author/funder. All rights reserved. No reuse allowed without permission.

bioRxiv preprint doi: https://doi.org/10.1101/2025.11.06.686738; this version posted November 7, 2025. The copyright holder for this preprint (which was not certified by peer review) is the author/funder. All rights reserved. No reuse allowed without permission.

We further interrogated whether the proportion of deconvoluted immune cells could segregate patients according to clinical groups. Our hierarchical clustering analysis

bioRxiv preprint doi: https://doi.org/10.1101/2025.11.06.686738; this version posted November 7, 2025. The copyright holder for this preprint (which was not certified by peer review) is the author/funder. All rights reserved. No reuse allowed without permission.

bioRxiv preprint doi: https://doi.org/10.1101/2025.11.06.686738; this version posted November 7, 2025. The copyright holder for this preprint (which was not certified by peer review) is the author/funder. All rights reserved. No reuse allowed without permission.

bioRxiv preprint doi: https://doi.org/10.1101/2025.11.06.686738; this version posted November 7, 2025. The copyright holder for this preprint (which was not certified by peer review) is the author/funder. All rights reserved. No reuse allowed without permission.

Most proteins, especially *N*-glycoproteins, were enriched for biological processes related to immune responses (Fig. 2C). In fact, tear fluid is essential for protecting the

bioRxiv preprint doi: https://doi.org/10.1101/2025.11.06.686738; this version posted November 7, 2025. The copyright holder for this preprint (which was not certified by peer review) is the author/funder. All rights reserved. No reuse allowed without permission.

bioRxiv preprint doi: https://doi.org/10.1101/2025.11.06.686738; this version posted November 7, 2025. The copyright holder for this preprint (which was not certified by peer review) is the author/funder. All rights reserved. No reuse allowed without permission.

For example, strong tissue expression of desmocollin 1 (DSC1) has been considered a predictor of epithelial dysplasia in oral lichen planus, a lesion that can also progress toward malignant transformation (43). In our data, DSC1 was upregulated in PVL and OSCC compared to OL. Of note, PVL is more prone to malignant progression than OL. Desmosomal proteins, such as desmoplakin (DSP), have been investigated in carcinogenesis because loss of cell adhesion is essential for tumor development and

bioRxiv preprint doi: https://doi.org/10.1101/2025.11.06.686738; this version posted November 7, 2025. The copyright holder for this preprint (which was not certified by peer review) is the author/funder. All rights reserved. No reuse allowed without permission.

##### Sample preparation for LC-MS/MS analysis

bioRxiv preprint doi: https://doi.org/10.1101/2025.11.06.686738; this version posted November 7, 2025. The copyright holder for this preprint (which was not certified by peer review) is the author/funder. All rights reserved. No reuse allowed without permission.

Quality evaluation of the LC–MS/MS runs was performed by measuring deviations in the retention times of three trypsin fragment peaks (m/z 421.7584, 2+; m/z 523.2855, 2+; and m/z 1106.0557, 2+). In addition, three iRT peptides were spiked into tear

bioRxiv preprint doi: https://doi.org/10.1101/2025.11.06.686738; this version posted November 7, 2025. The copyright holder for this preprint (which was not certified by peer review) is the author/funder. All rights reserved. No reuse allowed without permission.

The LC–MS/MS data were searched with the HCP workflow using Byonic v2.6.46 (Byos Software, Protein Metrics Inc., CA, USA) (23), with precursor and product ion mass tolerances of 10 ppm and 20 ppm, respectively. Trypsin-specific cleavages were

bioRxiv preprint doi: https://doi.org/10.1101/2025.11.06.686738; this version posted November 7, 2025. The copyright holder for this preprint (which was not certified by peer review) is the author/funder. All rights reserved. No reuse allowed without permission.

Data acquisition and fluorescence intensity analysis were carried out

bioRxiv preprint doi: https://doi.org/10.1101/2025.11.06.686738; this version posted November 7, 2025. The copyright holder for this preprint (which was not certified by peer review) is the author/funder. All rights reserved. No reuse allowed without permission.

**GO annotation of biological processes**

bioRxiv preprint doi: https://doi.org/10.1101/2025.11.06.686738; this version posted November 7, 2025. The copyright holder for this preprint (which was not certified by peer review) is the author/funder. All rights reserved. No reuse allowed without permission.

The clinical and pathological features included in the models are detailed in Table S5. Initially, a general model was constructed using features common to all clinical groups, including mean age (≥ vs. <45 years; ≥ vs. <60 years), gender (male/female), smoking habits (yes/no), alcohol consumption (yes/no), lesion location (tongue/other), and immune cell infiltrate based on histological evaluation

bioRxiv preprint doi: https://doi.org/10.1101/2025.11.06.686738; this version posted November 7, 2025. The copyright holder for this preprint (which was not certified by peer review) is the author/funder. All rights reserved. No reuse allowed without permission.

bioRxiv preprint doi: https://doi.org/10.1101/2025.11.06.686738; this version posted November 7, 2025. The copyright holder for this preprint (which was not certified by peer review) is the author/funder. All rights reserved. No reuse allowed without permission.


1. Bittar RF, Ferraro HP, Ribas MH, Lehn CN. Predictive factors of occult neck metastasis in patients with oral squamous cell carcinoma. Braz J Otorhinolaryngol. 2016 Sep-Oct;82(5):543-7. doi: 10.1016/j.bjorl.2015.09.005. Epub 2015 Dec 17. PMID: 26749457; PMCID: PMC9444684.
2. Wang Y, Yang T, Gan C, Wang K, Sun B, Wang M, Zhu F. Temporal and spatial patterns of recurrence in oral squamous cell carcinoma, a single-center retrospective cohort study in China. BMC Oral Health. 2023 Sep 19;23(1):679. doi: 10.1186/s12903-023-03204-7. PMID: 37726764; PMCID: PMC10510235.
3. Henry M, Rosberger Z, Bertrand L, Klassen C, Hier M, Zeitouni A, Kost K, Mlynarek A, Richardson K, Black M, MacDonald C, Zhang X, Chartier G, Frenkiel S. Prevalence and Risk Factors of Suicidal Ideation among Patients with Head and Neck Cancer: Longitudinal Study. Otolaryngol Head Neck Surg. 2018 Nov;159(5):843-852. doi: 10.1177/0194599818776873. Epub 2018 Jun 5. PMID: 29865939.
4. Piemonte ED, Gilligan GM, Garola F, Lazos JP, Panico RL, Normando AGC, Santos-Silva AR, Warnakulasuriya S. Differences among oral carcinomas arising de novo from those associated with oral potentially malignant disorders: a systematic review and meta-analysis. Oral Surg Oral Med Oral Pathol Oral Radiol. 2024 Jun;137(6):613-631. doi: 10.1016/j.oooo.2024.03.006. Epub 2024 Mar 19. PMID: 38609795.
5. Iocca O, Sollecito TP, Alawi F, Weinstein GS, Newman JG, De Virgilio A, et al. Potentially malignant disorders of the oral cavity and oral dysplasia: A systematic review and meta-analysis of malignant transformation rate by subtype. Head Neck. 2020 Mar;42(3):539-555.
6. Hvid-Jensen F, Pedersen L, Drewes AM, Sørensen HT, Funch-Jensen P. Incidence of adenocarcinoma among patients with Barrett's esophagus. N Engl J Med. 2011 Oct 13;365(15):1375-83. doi: 10.1056/NEJMoa1103042. PMID: 21995385.
7. Lodi G, Franchini R, Warnakulasuriya S, Varoni EM, Sardella A, Kerr AR, Carrassi A, MacDonald LC, Worthington HV. Interventions for treating oral leukoplakia to prevent oral

bioRxiv preprint doi: https://doi.org/10.1101/2025.11.06.686738; this version posted November 7, 2025. The copyright holder for this preprint (which was not certified by peer review) is the author/funder. All rights reserved. No reuse allowed without permission.

8. Odell E, Kujan O, Warnakulasuriya S, Sloan P. Oral epithelial dysplasia: Recognition, grading and clinical significance. Oral Dis. 2021 Nov;27(8):1947-1976. doi: 10.1111/odi.13993. Epub 2021 Sep 14. PMID: 34418233.
9. Thompson LDR, Fitzpatrick SG, Müller S, Eisenberg E, Upadhyaya JD, Lingen MW, Vigneswaran N, Woo SB, Bhattacharyya I, Bilodeau EA, Carlos R, Islam MN, Leon ME, Lewis JS Jr, Magliocca KR, Mani H, Mehrad M, Purgina B, Richardson M, Wenig BM, Cohen DM. Proliferative Verrucous Leukoplakia: An Expert Consensus Guideline for Standardized Assessment and Reporting. Head Neck Pathol. 2021 Jun;15(2):572-587. doi: 10.1007/s12105-020-01262-9. Epub 2021 Jan 7. PMID: 33415517; PMCID: PMC8134585.
10. Proaño-Haro A, Bagan L, Bagan JV. Recurrences following treatment of proliferative verrucous leukoplakia: A systematic review and meta-analysis. J Oral Pathol Med. 2021 Sep;50(8):820-828. doi: 10.1111/jop.13178. Epub 2021 Apr 13. PMID: 33765364.
11. Nakayasu ES, Gritsenko M, Piehowski PD, Gao Y, Orton DJ, Schepmoes AA, Fillmore TL, Frohnert BI, Rewers M, Krischer JP, Ansong C, Suchy-Dicey AM, Evans-Molina C, Qian WJ, Webb-Robertson BM, Metz TO. Tutorial: best practices and considerations for mass-spectrometry-based protein biomarker discovery and validation. Nat Protoc. 2021 Aug;16(8):3737-3760. doi: 10.1038/s41596-021-00566-6. Epub 2021 Jul 9. PMID: 34244696; PMCID: PMC8830262.
12. Sá JO, Trino LD, Oliveira AK, Lopes AFB, Granato DC, Normando AGC, Santos ES, Neves LX, Carnielli CM, Paes Leme AF. Proteomic approaches to assist in diagnosis and prognosis of oral cancer. Expert Rev Proteomics. 2021 Apr;18(4):261-284. doi: 10.1080/14789450.2021.1924685. Epub 2021 May 18. PMID: 33945368.
13. Surdu A, Foia LG, Luchian I, Trifan D, Tatarciuc MS, Scutariu MM, Ciupilan C, Budala DG. Saliva as a Diagnostic Tool for Systemic Diseases-A Narrative Review. Medicina (Kaunas). 2025 Jan 30;61(2):243. doi: 10.3390/medicina61020243. PMID: 40005360; PMCID: PMC11857487.
14. Tasoulas J, Patsouris E, Giaginis C, Theocharis S. Salivaomics for oral diseases biomarkers detection. Expert Rev Mol Diagn. 2016;16(3):285-95. doi: 10.1586/14737159.2016.1133296. Epub 2016 Jan 11. PMID: 26680995.
15. Nagler RM, Hershkovich O. Age-related changes in unstimulated salivary function and composition and its relations to medications and oral sensorial complaints. Aging Clin Exp Res. 2005 Oct;17(5):358-66. doi: 10.1007/BF03324623. PMID: 16392409.
16. Nicholson K, Liu W, Fitzpatrick D, Hardacre KA, Roberts S, Salerno J, Stranges S, Fortin M, Mangin D. Prevalence of multimorbidity and polypharmacy among adults and older adults: a

bioRxiv preprint doi: https://doi.org/10.1101/2025.11.06.686738; this version posted November 7, 2025. The copyright holder for this preprint (which was not certified by peer review) is the author/funder. All rights reserved. No reuse allowed without permission.

A comprehensive scoping review of methodological approaches and clinical applications of tear fluid biomarkers. Prog Retin Eye Res. 2025 May;106:101338. doi: 10.1016/j.preteyeres.2025.101338. Epub 2025 Feb 13. PMID: 39954936.
20. Fatima A, Sanyal S, Jha GK, Kaliki S, Pallavi R. The enigmatic world of tear extracellular vesicles (EVs)-exploring their role in ocular health and beyond. FEBS Lett. 2025 May;599(10):1346-1372. doi: 10.1002/1873-3468.70004. Epub 2025 Feb 17. PMID: 39961136.
21. Hu L, Liu X, Zheng Q, Chen W, Xu H, Li H, Luo J, Yang R, Mao X, Wang S, Chen T, Lee LP, Liu F. Interaction network of extracellular vesicles building universal analysis via eye tears: iNEBULA. Sci Adv. 2023 Mar 17;9(11):eadg1137. doi: 10.1126/sciadv.adg1137. Epub 2023 Mar 15. PMID: 36921051; PMCID: PMC10017052.
22. Runström G, Mann A, Tighe B. The fall and rise of tear albumin levels: a multifactorial phenomenon. Ocul Surf. 2013 Jul;11(3):165-80. doi: 10.1016/j.jtos.2013.03.001. Epub 2013 May 1. PMID: 23838018.
23. Bern M, Kil YJ, Becker C. Byonic: advanced peptide and protein identification software. Curr Protoc Bioinformatics. 2012 Dec;Chapter 13:13.20.1-13.20.14. doi: 10.1002/0471250953.bi1320s40. PMID: 23255153; PMCID: PMC3545648.
24. Escher C, Reiter L, MacLean B, Ossola R, Herzog F, Chilton J, MacCoss MJ, Rinner O. Using iRT, a normalized retention time for more targeted measurement of peptides. Proteomics. 2012 Apr;12(8):1111-21. doi: 10.1002/pmic.201100463. PMID: 22577012; PMCID: PMC3918884.
25. Nguyen-Khuong T, Everest-Dass AV, Kautto L, Zhao Z, Willcox MD, Packer NH. Glycomic characterization of basal tears and changes with diabetes and diabetic retinopathy. Glycobiology. 2015 Mar;25(3):269-83. doi: 10.1093/glycob/cwu108. Epub 2014 Oct 9. PMID: 25303961.
26. Ponzini E. Tear biomarkers. Adv Clin Chem. 2024;120:69-115. doi: 10.1016/bs.acc.2024.03.002. Epub 2024 Apr 11. PMID: 38762243.
27. Jones G, Lee TJ, Glass J, Rountree G, Ulrich L, Estes A, Sezer M, Zhi W, Sharma S, Sharma A. Comparison of Different Mass Spectrometry Workflows for the Proteomic Analysis of Tear

bioRxiv preprint doi: https://doi.org/10.1101/2025.11.06.686738; this version posted November 7, 2025. The copyright holder for this preprint (which was not certified by peer review) is the author/funder. All rights reserved. No reuse allowed without permission.

bioRxiv preprint doi: https://doi.org/10.1101/2025.11.06.686738; this version posted November 7, 2025. The copyright holder for this preprint (which was not certified by peer review) is the author/funder. All rights reserved. No reuse allowed without permission.

bioRxiv preprint doi: https://doi.org/10.1101/2025.11.06.686738; this version posted November 7, 2025. The copyright holder for this preprint (which was not certified by peer review) is the author/funder. All rights reserved. No reuse allowed without permission.

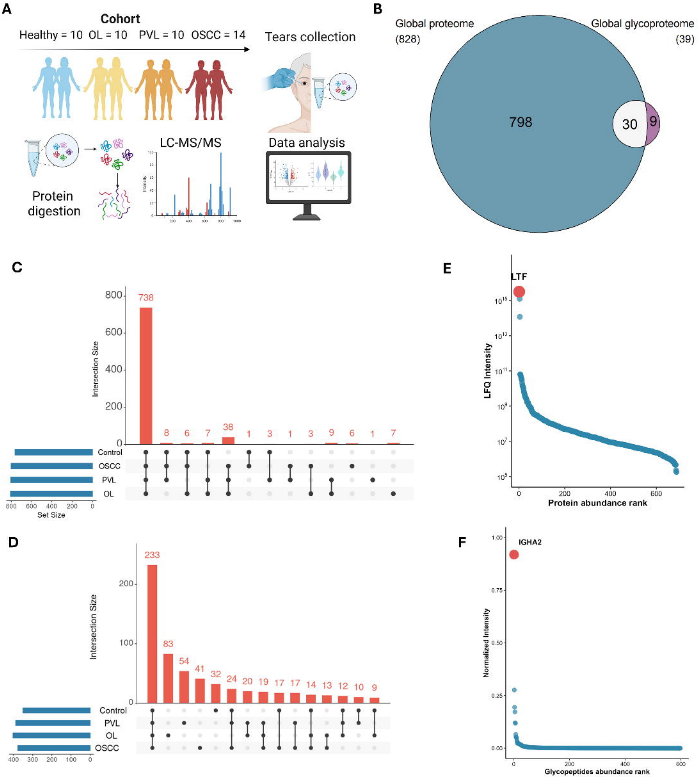
A
Cohort
Healthy = 10 OL = 10 PVL = 10 OSCC = 14
Tears collection
Protein digestion
LC-MS/MS
Data analysis
B
Global proteome (828)
Global glycoproteome (39)
798
30
9
C
Intersection Size
738
8
6
7
38
1
3
1
3
9
6
1
7
Control
OSCC
PVL
OL
Set Size
E
LTF
LFQ Intensity
Protein abundance rank
D
Intersection Size
233
83
54
41
32
24
20
19
17
17
14
13
12
10
9
Control
PVL
OL
OSCC
F
IGHA2
Normalized Intensity
Glycopeptides abundance rank

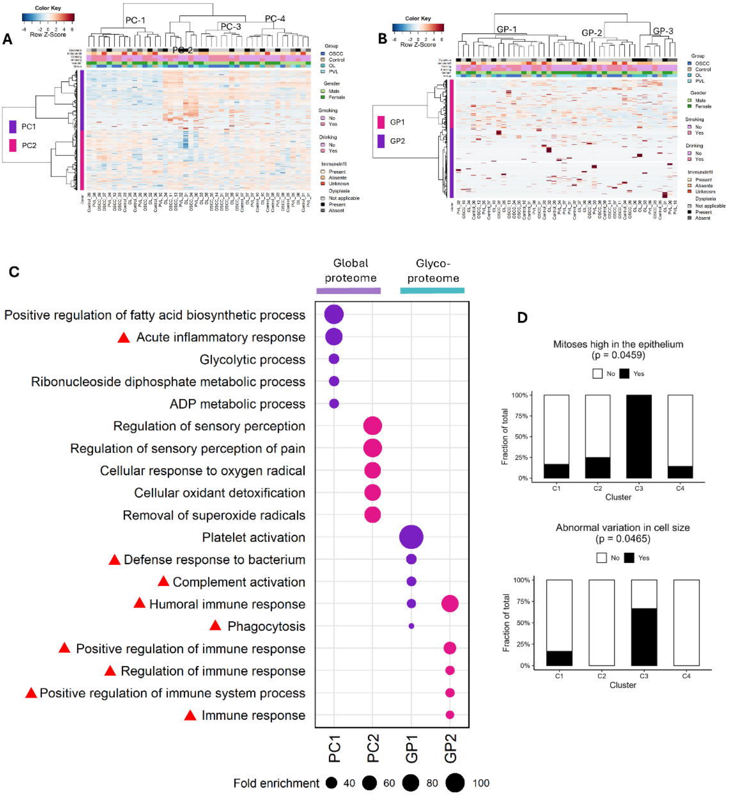
A
Color Key
Row Z-Score
PC-1
PC-2
PC-3
PC-4
PC1
PC2
Group
OSCC
Control
OL
PVL
Gender
Male
Female
Smoking
No
Yes
Drinking
No
Yes
ImmuneInfil
Present
Absente
Unknown
Dysplasia
Not applicable
Present
Absent
B
Color Key
Row Z-Score
GP-1
GP-2
GP-3
GP1
GP2
C
Global proteome
Glyco-proteome
Positive regulation of fatty acid biosynthetic process
Acute inflammatory response
Glycolytic process
Ribonucleoside diphosphate metabolic process
ADP metabolic process
Regulation of sensory perception
Regulation of sensory perception of pain
Cellular response to oxygen radical
Cellular oxidant detoxification
Removal of superoxide radicals
Platelet activation
Defense response to bacterium
Complement activation
Humoral immune response
Phagocytosis
Positive regulation of immune response
Regulation of immune response
Positive regulation of immune system process
Immune response
PC1
PC2
GP1
GP2
Fold enrichment
40
60
80
100
D
Mitoses high in the epithelium
(p = 0.0459)
No
Yes
Fraction of total
100%
75%
50%
25%
0%
C1
C2
C3
C4
Cluster
Abnormal variation in cell size
(p = 0.0465)
No
Yes
Fraction of total
100%
75%
50%
25%
0%
C1
C2
C3
C4
Cluster

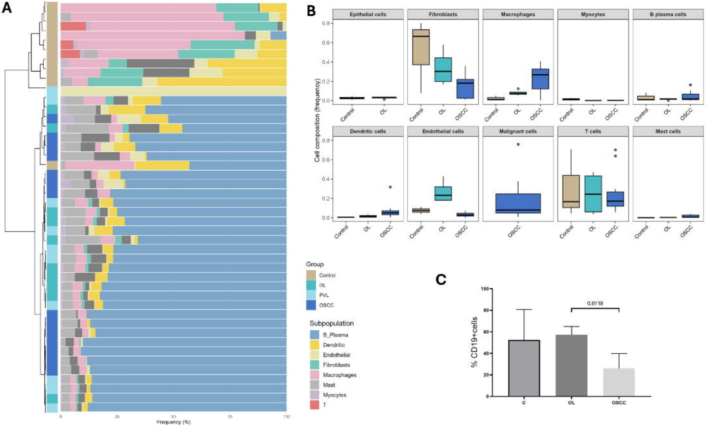
A
0
25
50
75
100
Frequency (%)
Group
Control
OL
PVL
OSCC
Subpopulation
B_Plasma
Dendritic
Endothelial
Fibroblasts
Macrophages
Mast
Myocytes
T
B
Cell composition (frequency)
Epithelial cells
Fibroblasts
Macrophages
Myocytes
B plasma cells
Dendritic cells
Endothelial cells
Malignant cells
T cells
Mast cells
0.0
0.2
0.4
0.6
0.8
Control
OL
OSCC
C
% CD19+cells
0
20
40
60
80
100
0.0118
C
OL
OSCC

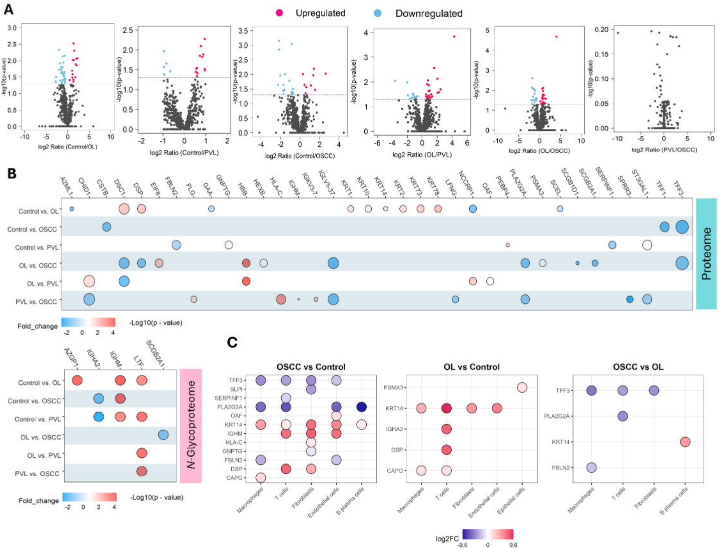

A
Upregulated
Downregulated
log2 Ratio (Control/OL)
log2 Ratio (Control/PVL)
log2 Ratio (Control/OSCC)
log2 Ratio (OL/PVL)
log2 Ratio (OL/OSCC)
log2 Ratio (PVL/OSCC)
-log10(p-value)
B
Control vs. OL
Control vs. OSCC
Control vs. PVL
OL vs. OSCC
OL vs. PVL
PVL vs. OSCC
Proteome
Fold_change
-Log10(p - value)
N-Glycoproteome
C
OSCC vs Control
OL vs Control
OSCC vs OL
Macrophages
T cells
Fibroblasts
Endothelial cells
B plasma cells
Epithelial cells
log2FC

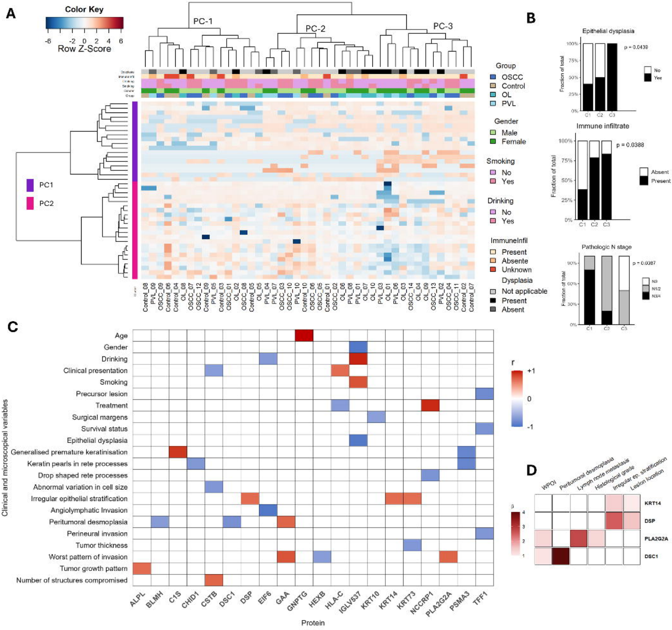
A
Color Key
-6
-2 0 2 4 6
Row Z-Score
PC-1
PC-2
PC-3
PC1
PC2
Group
OSCC
Control
OL
PVL
Gender
Male
Female
Smoking
No
Yes
Drinking
No
Yes
ImmuneInfil
Present
Absente
Unknown
Dysplasia
Not applicable
Present
Absent
Control_08
PVL_09
OSCC_09
Control_06
Control_04
OL_08
OSCC_07
OSCC_12
Control_09
PVL_05
Control_03
OSCC_01
OL_02
OSCC_08
Control_05
OL_05
PVL_04
PVL_07
OSCC_03
OSCC_10
PVL_10
Control_10
OSCC_06
OSCC_05
Control_01
OSCC_02
OL_06
PVL_08
PVL_01
OL_07
OL_10
PVL_03
OL_01
PVL_06
OL_03
OSCC_14
OL_04
OL_09
OSCC_13
PVL_02
OSCC_04
OSCC_11
Control_02
Control_07
B
Epithelial dysplasia
p = 0.0438
No
Yes
Fraction of total
C1
C2
C3
Immune infiltrate
p = 0.0388
Absent
Present
Pathologic N stage
p = 0.0387
N0
N1/2
N3/4
C
Age
Gender
Drinking
Clinical presentation
Smoking
Precursor lesion
Treatment
Surgical margens
Survival status
Epithelial dysplasia
Generalised premature keratinisation
Keratin pearls in rete processes
Drop shaped rete processes
Abnormal variation in cell size
Irregular epithelial stratification
Angiolymphatic Invasion
Peritumoral desmoplasia
Perineural invasion
Tumor thickness
Worst pattern of invasion
Tumor growth pattern
Number of structures compromised
Clinical and microscopical variables
ALPL
BLMH
C1S
CHID1
CSTB
DSC1
DSP
EIF6
GAA
GNPTG
HEXB
HLA-C
IGLV537
KRT10
KRT14
KRT73
NCCRP1
PLA2G2A
PSMA3
TFF1
Protein
r
+1
0
-1
D
WPOI
Peritumoral desmoplasia
Lymph node metastasis
Histological grade
Irregular ep. stratification
Lesion location
KRT14
DSP
PLA2G2A
DSC1
